# Sequence modeling and design from molecular to genome scale with Evo

**DOI:** 10.1101/2024.02.27.582234

**Authors:** Eric Nguyen, Michael Poli, Matthew G. Durrant, Armin W. Thomas, Brian Kang, Jeremy Sullivan, Madelena Y. Ng, Ashley Lewis, Aman Patel, Aaron Lou, Stefano Ermon, Stephen A. Baccus, Tina Hernandez-Boussard, Christopher Ré, Patrick D. Hsu, Brian L. Hie

## Abstract

The genome is a sequence that completely encodes the DNA, RNA, and proteins that orchestrate the function of a whole organism. Advances in machine learning combined with massive datasets of whole genomes could enable a biological foundation model that accelerates the mechanistic understanding and generative design of complex molecular interactions. We report Evo, a genomic foundation model that enables prediction and generation tasks from the molecular to genome scale. Using an architecture based on advances in deep signal processing, we scale Evo to 7 billion parameters with a context length of 131 kilobases (kb) at single-nucleotide, byte resolution. Trained on 2.7M prokaryotic and phage genomes, Evo can generalize across the three fundamental modalities of the central dogma of molecular biology to perform zero-shot function prediction that is competitive with, or outperforms, leading domain-specific language models. Evo also excels at multi-element generation tasks, which we demonstrate by generating synthetic CRISPR-Cas molecular complexes and entire transposable systems for the first time. Using information learned over whole genomes, Evo can also predict gene essentiality at nucleotide resolution and can generate coding-rich sequences up to 650 kb in length, orders of magnitude longer than previous methods. Advances in multi-modal and multiscale learning with Evo provides a promising path toward improving our understanding and control of biology across multiple levels of complexity.

## 1. Introduction

DNA is the fundamental layer of biological information that is responsible for transmitting the results of evolution across generations of life (Morgan, 1910; Watson and Crick, 1953; Nirenberg and Matthaei, 1961). Evolutionary variation in genome sequences is a reflection of adaptation and selection for biological function at the phenotypic level (Dobzhansky, 1951). Rapid advances in DNA sequencing technologies have enabled the systematic mapping of this evolutionary diversity at the whole-genome scale.

A machine that learns this breadth of information across genomes could model the function of DNA, RNA, and proteins, as well as their diverse interactions that orchestrate complex biological functions, mediate disease, or create a complete organism. Modern machine learning algorithms combined with massive datasets of genomic sequences could enable a general biological foundation model that learns the intrinsic logic of whole genomes.

However, current efforts to model molecular biology with machine learning have been focused on creating modality-specific models that are specialized to proteins, regulatory DNA, or RNA (Jumper et al., 2021; Rives et al., 2021; Avsec et al., 2021; Theodoris et al., 2023). In addition, generative applications in biology have been limited to the design of single molecules, simple complexes (Watson et al., 2023; Madani et al., 2023 Ingraham et al., 2023), or short DNA sequences (DaSilva et al., 2024; Lal et al., 2024). In contrast, complex biological processes, such as gene regulation, CRISPR immunity, or genetic transposition, rely on many interactions involving molecules across multiple modalities.

A DNA model that unifies information across the molecular, systems, and genome scale could learn from large genomic regions to capture systems-wide interactions and enable the design of more sophisticated biological functions. By operating at single-nucleotide resolution, this model would be able to incorporate the evolutionary effects of sequence variation, such as individual single-nucleotide mutations that completely alter organism function.

Inspired by the recent success of large language models, many contemporary approaches leverage autoregressive Transformers to model biological sequences and to capture these system-wide interactions. However, existing attempts to model DNA as a language (Zvyagin et al., 2023; Dalla-Torre et al., 2023; Zhou et al., 2023) are limited by the prevailing dense Transformer architecture, which incurs high computational cost as input sequence lengths grow relative to model width (scaling quadratically) and generally underperforms at single-nucleotide or byte-level resolution (compared to models trained at coarser resolutions) (Tay et al., 2021). Recent algorithmic advances in extending context length of attention-based models (Chen et al., 2023; Liu et al., 2023) have similar resolution limitations. As a result, Transformer-based DNA models are constrained to short context lengths and use schemes that aggregate nucleotides into the basic units of language models, called tokens, thereby sacrificing single-nucleotide resolution (Zvyagin et al., 2023; Dalla-Torre et al., 2023; Fishman et al., 2023; Ji et al., 2021; Hwang et al., 2023).

Here, we present Evo, a 7 billion parameter genomic foundation model trained to generate DNA sequences at whole-genome scale. Evo uses a context length of 131k tokens and is based on the StripedHyena architecture (Poli et al., 2023b), which hybridizes attention and data-controlled convolutional operators to efficiently process and recall patterns in long sequences. Evo is trained on a prokaryotic whole-genome dataset consisting of 300 billion nucleotides and uses a byte-level, single-nucleotide tokenizer.

We demonstrate that Evo can be used in both prediction and generation tasks at the molecular, systems, and genome scale. In zero-shot evaluations, Evo is competitive with state-of-the-art protein language models at predicting the fitness effects of mutations on *E. coli* proteins, outperforms specialized RNA language models in predicting fitness effects of mutations on noncoding RNAs, and predicts the combinations of prokaryotic promoter-ribosome binding site (RBS) pairs that lead to active gene expression from regulatory sequence alone. Moving beyond single molecules and short sequences, Evo learns the co-evolutionary linkage of coding and noncoding sequences in order to design synthetic multi-component biological systems including CRISPR-Cas systems and transposable elements. At the whole-genome scale, Evo can predict essential genes in bacteria or bacteriophages without any supervision. We also use Evo to generate sequences over 650 kilobases (kb) with plausible genomic coding architecture, a scale that is orders of magnitude greater than previous methods (DaSilva et al., 2024; Lal et al., 2024; Watson et al., 2023). Taken together, Evo establishes a foundational paradigm for predictive and generative biological sequence modeling (Figure 1A). Further development of Evo will enable a deeper mechanistic understanding of biology and accelerate our ability to engineer life.

**Figure 1.**
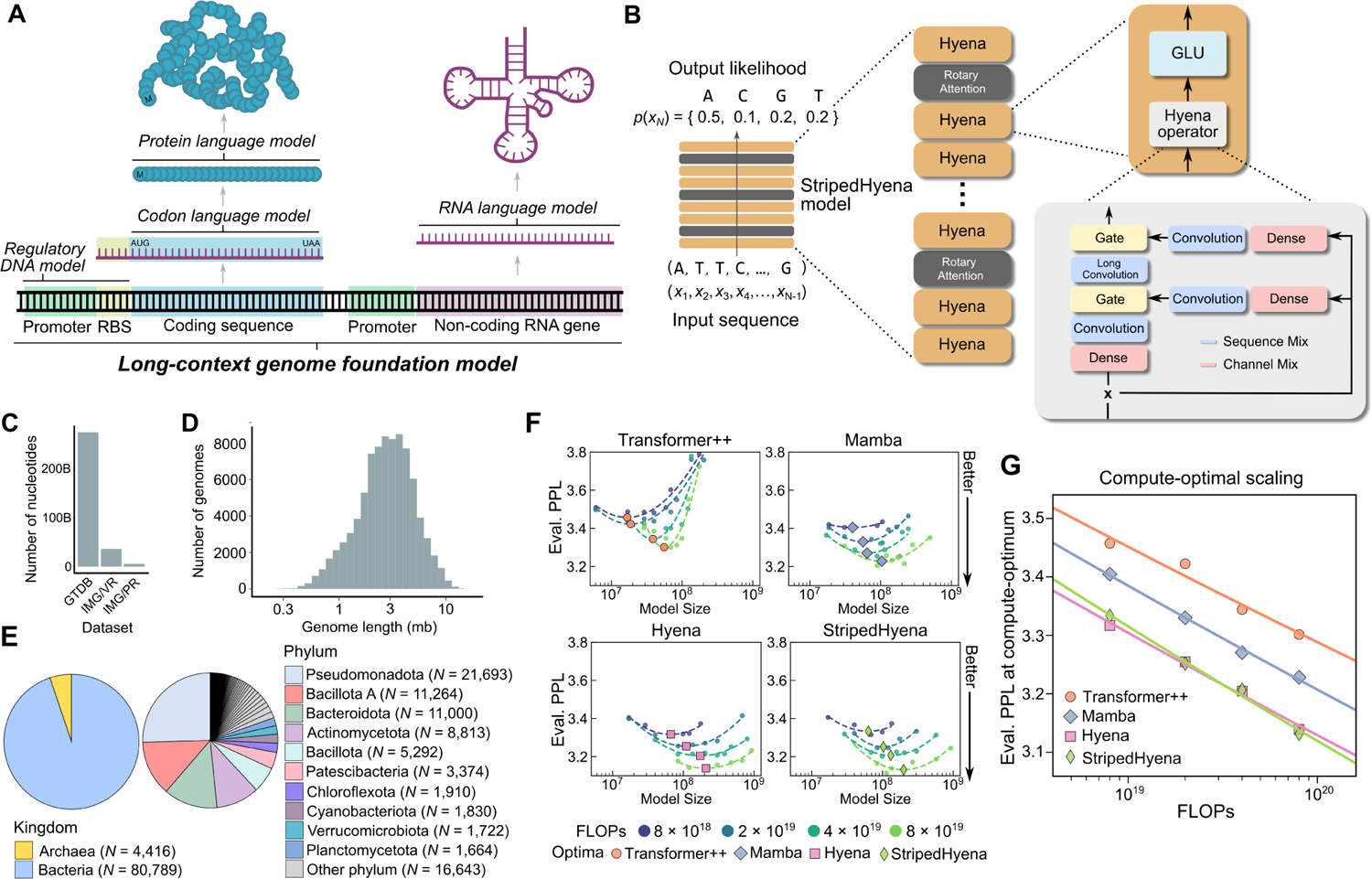
Pretraining a genomic foundation model across prokaryotic life. (**A**) A model of genome sequences at single-nucleotide resolution could learn all of the information encoded in regulatory DNA and in the sequences of the other modalities within the central dogma (proteins, coding RNA, and noncoding RNA). Even further, it could learning covariation involving multiple genes and regulatory elements. The status of DNA as the fundamental layer of biological information makes it a productive modality at which to develop a biological foundation model. (**B**) A model that predicts the likelihood of the next token given a sequence of tokens, referred to as autoregressive modeling, can learn complex patterns underlying DNA sequences. StripedHyena is a deep signal processing architecture for long sequences, obtained by hybridizing attention and hyena operators. (**C**) We pretrained Evo, a 7B parameter model with the StripedHyena architecture, on bacterial genome sequences from GTDB and IMG/PR and viral sequences from IMG/VR, excluding sequences from viruses that infect eukaryotic hosts. (**D**) A histogram depicting the sequence length of the genomes in GTDB. mb: megabases. (**E**) Pie charts depicting the taxonomic makeup of GTDB based on the kingdom (left) and phylum (right). (**F**) Results from a first-of-its-kind scaling laws analysis for large-scale DNA pretraining. Models improve monotonically with scale, with significant differences between architectures. Eval. PPL: evaluation perplexity. (**G**) To determine optimal architecture and scaling for Evo, we compared scaling rates of different models pretrained on the compute–optimal frontier, i.e., with optimal allocation of compute between dataset size and model size. Eval. PPL: evaluation perplexity. FLOPs: Floating point operations.

## 2. Results

### 2.1. Modeling long sequences at nucleotide resolution with the StripedHyena architecture

Evo is a genomic foundation model with 7B parameters trained with a context length of up to 131k tokens, using single-nucleotide, byte-level tokenization. To model long sequences at nucleotide resolution efficiently, which we demonstrate by generating sequences over 650k tokens, we leveraged the StripedHyena architecture (Poli et al., 2023b) (Figure 1B) that builds on emerging techniques in deep signal processing (Li et al., 2020; Gu et al., 2021; Orvieto et al., 2023; Massaroli et al., 2024). The model is a hybrid of 29 layers of data-controlled convolutional operators (hyena layers) interleaved with 3 layers (10%) of multi-head attention equipped with rotary position embeddings (RoPE) (Su et al., 2024) (**Methods**).

Model hybridization, first proposed to address shortcomings of state-space models (Ma et al., 2022; Fu et al., 2022; Pilault et al., 2024) has recently been shown to improve scaling performance on language modeling of both standalone Hyena and Transformer architectures (Poli et al., 2023b). StripedHyena is designed to benefit from the specialization of each of its layer types, with hyena layers implementing the bulk of the computation required for sequence processing and attention layers supplementing the ability to recall information from the context of an input.

Hyena layers process sequences in an input-dependent manner via compositions of short and long convolution filters (Figure 1B), making the layer especially effective at filtering noisy patterns that can occur in DNA and at aggregating individual nucleotides into motifs. Compared to HyenaDNA (Nguyen et al., 2023), a previous generation of DNA models leveraging a Hyena architecture (Poli et al., 2023a), Evo is based on an improved hybrid design and scaled to 1000× larger model size and 100× more data.

### 2.2. Training Evo at scale on OpenGenome

We compiled a large genome dataset called OpenGenome (**Methods**) with over 80, 000 bacterial and archaeal genomes, and millions of predicted prokaryotic phage and plasmid sequences, covering 300B nucleotide tokens (Figures 1C and **S1**) (Parks et al., 2022; Camargo et al., 2023, 2024). For safety considerations, we excluded viral genomes that infect eukaryotic hosts. Like most language models, Evo is pretrained via a next-token prediction objective on raw genome sequences with no explicit supervision or annotations. In order to predict the next token given a sequence of tokens, the model must learn the distribution of the genome data and be aware of the biological sequence motifs found in the collected genomes. Pretraining involves 2 stages: the first stage uses a context length of 8k tokens, while the second context extension stage uses 131k tokens as context. Depending on the downstream task, we select a base model from one of the two stages to finetune on smaller datasets of interest for generation.

### 2.3. StripedHyena demonstrates favorable scaling laws on DNA sequence data

Aiding our model design, we performed the first scaling laws analysis (to our knowledge) for DNA sequence modeling. The main objective of this type of analysis is to determine the relationship between training, architectural details, and performance metrics via a systematic experimental protocol (Hoffmann et al., 2022; Kaplan et al., 2020). Once a set of scaling laws is obtained, it can then be used as a guide to optimally scale training to larger models and datasets.

Here, we compare different classes of architectures via a compute-optimal protocol, aimed at evaluating results on the *compute-optimal frontier* (**Methods**). We trained over 300 models across four architectures: Transformer++, Mamba, Hyena, and StripedHyena. Transformer++ is a state-of-the-art Transformer, and Mamba is a modern architecture using data-controlled state-space models (Gu and Dao, 2023).

We found Transformer++ to yield significantly worse perplexity (a measure of next token prediction quality) at all compute budgets (Figures 1G), a symptom of the inefficiency of the architecture at the byte resolution. State-space and deep signal processing architectures are observed to improve on the scaling rate over Transformer++, with Hyena and StripedHyena resulting in the best scaling rate. We observed stable training for StripedHyena throughout all the studied model sizes and learning rates during the scaling analysis.

We also compare architecture performance outside the compute-optimal frontier, namely with allocations of the computational budget that may be suboptimal. Performance outside the compute-optimal frontier is important in practice, as most models (including Evo) are trained for more tokens than recommended by compute-optimal scaling laws. We estimate 250 billion to be the compute-optimal number of tokens for Evo 7B given the FLOP budget, meaning the model was trained at a 17% offset from the compute-optimal model size during the initial 8192 sequence length pretraining phase of 300 billion tokens. Both Transformer++ and Mamba experienced numerical instability during training, and suffered from a sharper performance degradation of the scaling rate outside the compute-optimal frontier, in contrast to StripedHyena (further analysis in **Figure S3**). These findings motivate the choice of StripedHyena as the architecture for Evo.

### 2.4. Evo performs zero-shot function prediction across DNA, RNA, and protein modalities

#### 2.4.1. Predicting mutational effects on protein function

Beyond evaluating perplexity, we investigated the model’s zero-shot performance on biologically relevant downstream tasks. For example, language models specifically trained on large corpuses of protein sequences or nucleotide coding sequences have demonstrated an impressive ability to predict mutational effects on protein function (Meier et al., 2021; Notin et al., 2022; Benegas et al., 2023) without any task-specific finetuning or supervision. Because Evo is trained on long genomic sequences that contain protein coding sequences, we tested whether the model would also learn the protein language well enough to perform zero-shot protein function prediction.

Following work in evaluation of protein language models, we leveraged deep mutational scanning (DMS) studies which introduce an exhaustive set of mutations to a protein coding sequence and then experimentally measure the effects of these mutations on various definitions of fitness (fitness is a study-specific metric quantifying how well a protein performs a certain function) (Notin et al., 2022, 2023; Livesey and Marsh, 2023). The language-model likelihood or pseudolikelihood (**Methods**) of the amino acid sequence is used to predict the experimental fitness score (Figure 2A). To adapt this task to nucleotide sequences, we use the wild-type coding sequence and nucleotide mutations reported in the original DMS studies (**Methods**).

**Figure 2.**
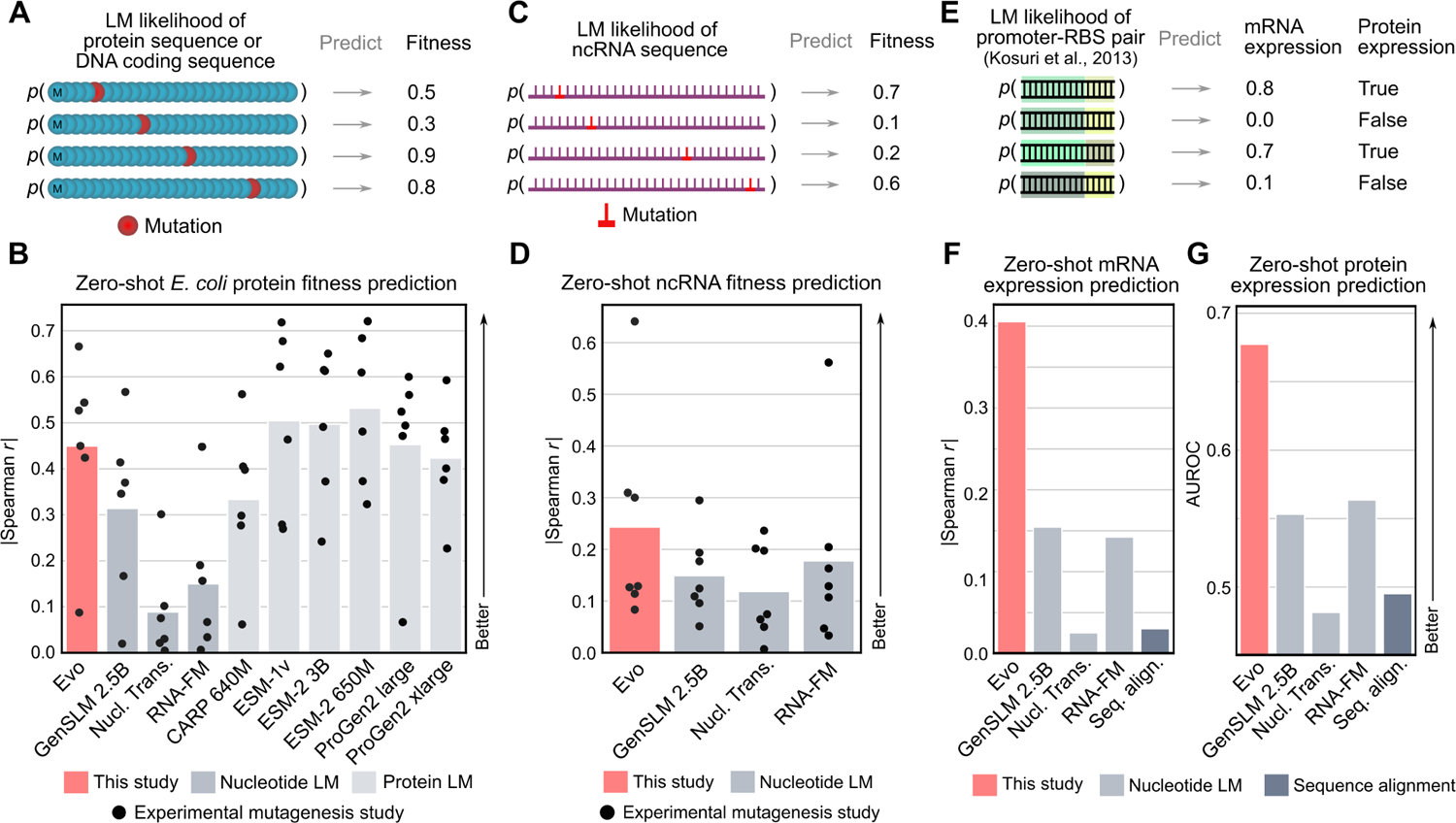
Evo performs zero-shot function prediction for proteins, non-coding RNAs, and regulatory DNA. (**A**) We obtained deep mutational scanning (DMS) datasets in which many mutations are made to a protein and a corresponding fitness score is experimentally measured for each protein variant. On the same set of mutated sequences, we compute its likelihood (or pseudolikelihood) under a protein language model or a nucleotide language model (LM). We then correlated these likelihoods with the experimental fitness measurements and used the strength of the correlation to measure the performance of zero-shot function prediction. (**B**) Evo has comparable predictive performance, measured via Spearman correlation, to state-of-the-art protein language models and higher performance than nucleotide language models. Bar height indicates the mean; each dot indicates a different DMS study. LM: language model; Nucl. Trans.: Nucleotide Transformer. (**C**) We obtained datasets in which many mutations are made to a ncRNA and a corresponding fitness score is experimentally measured. Predictive performance is measured as in the method described in (**A**). (**D**) Evo exhibits higher performance than nucleotide language models at predicting mutational effects on ncRNA function. Bar height indicates the mean; each dot indicates a different DMS study. LM: language model; Nucl. Trans.: Nucleotide Transformer. (**E**) We obtained a dataset in which Kosuri et al. (2013) measured mRNA and protein expression of a gene downstream of ∼12k promoter-RBS pairs in *E. coli*. For each promoter-RBS pair, we computed the likelihood of the sequence under a language model or a score indicating the frequency with which a promoter-RBS pair is observed in bacterial genomes. (**F**, **G**) Evo has higher predictive performance of mRNA and protein expression compared to nucleotide language models and to methods for computing the frequency of promoter-RBS pairs based on sequence alignment (“Seq. align.”). Bar height indicates the mean; each dot indicates a different DMS study. LM: language model; Nucl. Trans.: Nucleotide Transformer.

When we evaluated Evo’s zero-shot ability to predict mutational effects on protein function using DMS datasets of *E. coli* proteins, we found that it outperformed all other nucleotide models tested (Figure 2B), including GenSLM (Zvyagin et al., 2023), a model explicitly trained only on coding sequences with a codon vocabulary (Figure 1A). Evo also reaches competitive performance with leading protein-specific language models (Yang et al., 2024; Meier et al., 2021; Lin et al., 2023; Madani et al., 2023) at this task (Figure 2B). Previous work has shown that improvement beyond this performance range is very difficult for protein language models with self-supervised pretraining alone (Li et al., 2024), indicating that Evo is already competitive with state-of-the-art protein language modeling on bacterial proteins. Notably, Evo is trained on long-context genomic sequences without any explicit coding sequence annotations. On DMS datasets of human proteins, Evo is unable to predict mutational effects on fitness (**Figure S6A**), most likely because the pretraining dataset only contains prokaryotic sequences. However, we observed a strong association between language-model perplexity on the wildtype sequence and fitness prediction performance (**Figure S6B**), indicating that additional finetuning or future pretraining on mammalian coding sequences could improve Evo’s performance beyond bacterial proteins.

#### 2.4.2. Predicting mutational effects on ncRNA function

Next, we tested whether the same pretrained model could learn functional information about noncoding RNAs (ncRNA), such as tRNAs, rRNAs, and ribozymes. ncRNAs are encoded in the genome in a similar manner to proteins and they serve a variety of essential functions, including in protein synthesis and gene regulation. We collected ncRNA DMS (**Methods**), which are conceptually similar to protein DMS datasets but where mutations are made to the ncRNA sequence instead. We likewise evaluated Evo’s ability to perform zero-shot ncRNA fitness prediction using the results of experimental ncRNA DMS studies as the ground truth score (Figure 2C).

We found that Evo again outperforms all other tested nucleotide language models at this task, including RNA-FM (Chen et al., 2022), an RNA language model that is explicitly trained on ncRNA sequences (Figure 2D). We observed especially strong predictive performance on a study that measured the effects of mutations to the 5S ribosomal RNA on the growth rate of *E. coli* (Spearman *r* = 0.64, two-sided *t*-distributed *P* = 7.3×10^−4^) (Zhang et al., 2009). Together with our results for protein sequences, these results indicate that Evo is able to learn from its prokaryotic genome training data to predict functional properties across different molecular modalities.

#### 2.4.3. Predicting gene expression from regulatory DNA

Given that Evo is also trained on prokaryotic regulatory DNA sequences in addition to sequences that encode proteins or ncRNA, we investigated whether it is able to learn aspects of DNA regulatory grammar. To this end, we leveraged a dataset in which Kosuri et al. (2013) constructed approximately 12k combinations of common promoters and ribosome binding sites (RBSs) and measured the corresponding mRNA and protein expression of a reporter gene for each promoter-RBS pair in *E. coli* (Figure 2E). We find that the model likelihood scores that Evo assigns to promoter-RBS sequences is significantly correlated with mRNA expression (Spearman *r* = 0.41, two-sided *t*-distributed *P* < 1 × 10^−5^) (Figure 2F) and predictive of binarized protein expression (area under the receiver operating characteristic curve [AUROC] = 0.68, permutation-based *P* < 1×10^−5^) (Figure 2G) (**Methods**). Evo’s predictive performance is also substantially higher than those of other nucleotide language models, though none of these baseline language models have been trained on datasets containing regulatory elements.

To predict gene expression from the promoter-RBS sequence alone, Evo most likely uses its knowledge of natural regulatory sequences learned during pretraining, analogous to how protein language models can predict functional changes based on natural variation in protein sequences (Meier et al., 2021; Notin et al., 2023). To this end, as an additional benchmark, we aligned the promoter-RBS pairs to our model’s pretraining dataset of prokaryotic sequences. While we hypothesized that the number or the strength of these alignments would be predictive of gene expression, we found that computing these alignments using standard bioinformatic tools resulted in poor or nonexistent sequence matches (**Methods**). Of all techniques we attempted, none were predictive of gene expression (Figures 2F and **2G**), indicating that Evo can distill non-obvious functional information of regulatory DNA directly from large genomic sequence databases.

Overall, we show how a single foundation model of prokaryotic genomes can perform tasks that have previously been accomplished by different, domain-specific models (protein language models, RNA language models, and regulatory DNA models). Despite being trained on long genomic crops without explicit sequence annotations, Evo still demonstrates an understanding of the constitutive protein-coding sequences, ncRNA sequences, and regulatory elements.

### 2.5. Generative design of CRISPR-Cas molecular complexes

Next, we reasoned that Evo should be able to generate functional complexes that involve interactions between distinct molecular modalities. In prokaryotes, functionally related genes are generally located next to each other on the linear genome sequence. Because Evo learns covariation patterns involving any genetic element within its context window, the model should understand interactions between encoded protein and ncRNA molecules. To demonstrate this capability, we finetuned Evo on a dataset of genomic loci containing CRISPR-Cas sequences: molecular machines that consist of one or more protein components and one or more ncRNA components that, together, direct adaptive immunity against viral infection (Wang et al., 2022).

The DNA-targeting Cas9 nuclease is typically encoded within 3,000 to 4,800 bp of coding sequence and found in close genomic proximity to its cognate CRISPR array (Hsu et al., 2014). Transcription from the CRISPR array generates non-coding CRISPR RNA (crRNA) molecules that are bound by the Cas protein to generate a functional defense complex that is required for sequence-specific DNA-targeting (Figure 3A). For Cas9 in particular, a second trans-activating CRISPR RNA (tracrRNA) forms a duplex with the crRNA to create a full guide RNA (gRNA). Diverse families of CRISPR-Cas systems are found throughout bacterial and archaeal life, such as Cas12- or Cas13-based systems that target DNA and RNA, respectively (Koonin and Makarova, 2019).

**Figure 3.**
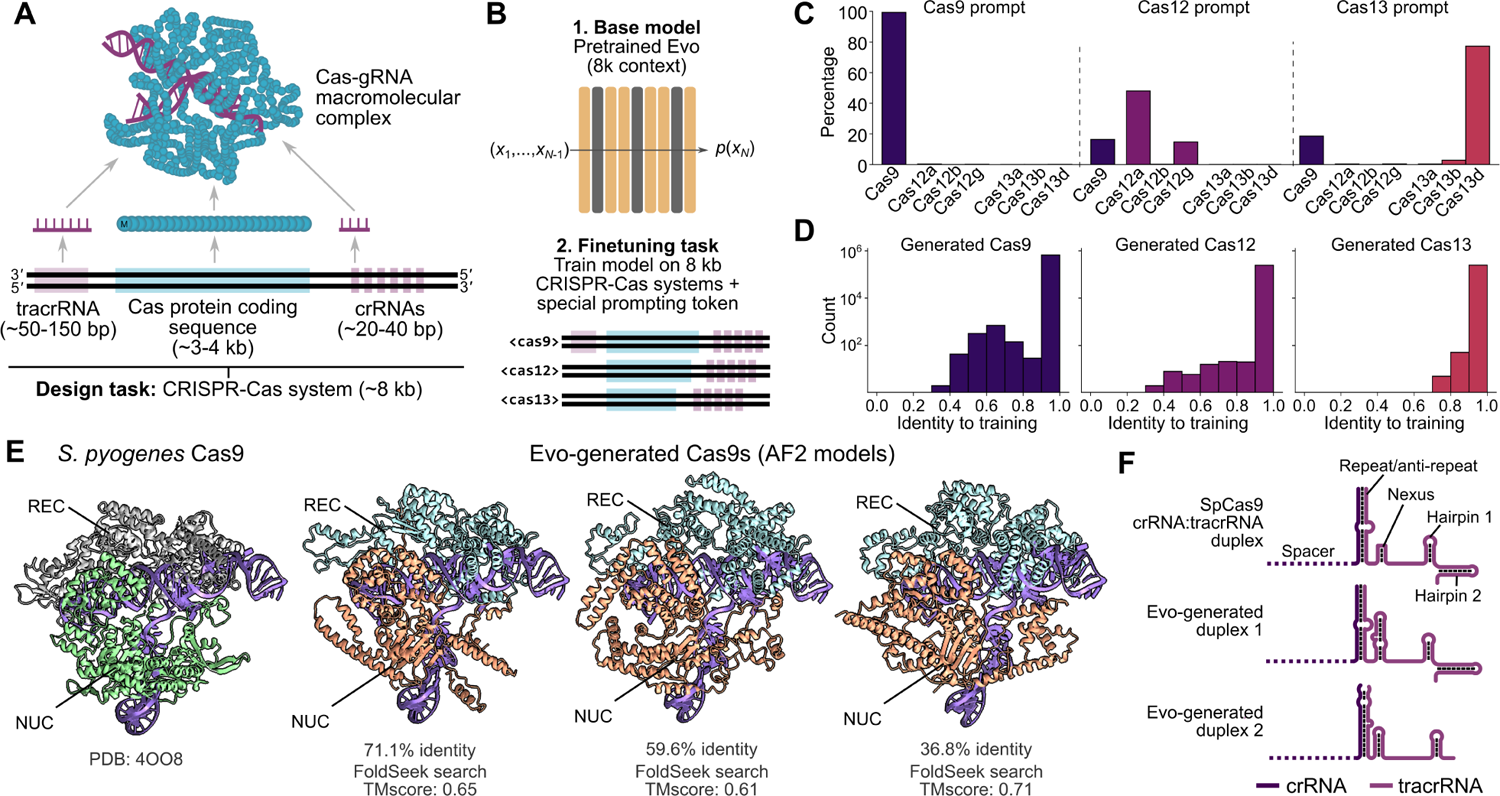
Finetuning on CRISPR-Cas sequences enables generative design of protein-RNA complexes. (A) CRISPR-Cas defense nucleases comprise a large macromolecular complex involving an effector protein bound to a noncoding guide RNA (gRNA) that is derived from a CRISPR RNA (crRNA). For some CRISPR types, a trans-activating CRISPR RNA (tracrRNA) is combined with the crRNA to create the final gRNA. Our design task is to produce sequences that contain these protein and noncoding RNA components. (B) We finetuned Evo, following its initial 8k pretraining phase, on 8 kb-length genomic sequences containing CRISPR-Cas systems. During finetuning, we prepended a special conditioning token (“cas9,” “cas12”, or “cas13”) to the beginning of each sequence, indicating the general type of Cas protein encoded in the sequence. (C) A prompting token enables controllability over Evo generations. When prompting with the token for a given type of Cas protein, the most common Cas protein found in the resulting generated sequences corresponds to that token prompt (for example, prompting with a “cas9” token typically produces Cas9 sequences). (D) Histograms representing the distribution of percentage identity of a generated Cas protein sequence to any Cas protein sequence in the training dataset. This distribution is computed across sampling runs involving all three prompts. (E) Representative generations of Cas proteins alongside the *S. pyogenes* Cas9 crystal structure (PDB: 4OO8). AF2: AlphaFold2. REC: recognition lobe. NUC: nuclease lobe. (F) Example generations of crRNA:tracrRNA duplexes alongside a canonical *S. pyogenes* Cas9 duplex.

In the finetuning step, we trained the model on 82,430 CRISPR-Cas loci extracted from public metagenomic and genomic sequences, adding special prompt tokens for Cas9, Cas12, and Cas13 that were prepended to the beginning of each training sequence (Figure 3B). During sampling, these tokens allow us to guide generation of a specific CRISPR-Cas system type by prompting with the corresponding special token. Strikingly, sampling 8 kb sequences using each of the three Cas token prompts resulted in coherent generations. Depending on the prompt token used, 15-45% of generations contained Cas coding sequences as long as 5kb as detected by Cas subtype profile HMMs (**Methods**). We also observed that prompting with a specific Cas subtype token typically produced a sample with the expected subtype, demonstrating that Evo can be tuned to generate sequences with both proteins of interest as well as associated non-coding elements such as CRISPR arrays (Figure 3C). Sequence alignment with the training dataset revealed that Evo is capable of highly unique Cas protein generations, as some of the predicted ORFs exhibited less than 40% protein sequence identity to their respective closest match (Figures 3D and **S7**).

To evaluate the quality of Cas generation with Evo, we focused on Cas9 generations and evaluated AlphaFold2 structure predictions of the sampled Cas9 coding sequence and non-coding RNA complexes against experimentally determined structures of the *Streptococcus pyogenes* Cas9 protein and its tracrRNA:crRNA duplex. Selected structure predictions of sampled Cas9 sequences show that even low-identity generations bear resemblance to natural Cas9 structures in key domains such as the RuvC nuclease and protospacer adjacent motif (PAM)-interacting domains (Figure 3E). Similarly, Evo-generated crRNA:tracrRNA duplexes form predicted RNA secondary structures resembling the canonical crRNA:tracrRNA duplexes found in naturally occurring Cas9 systems (Figure 3F) (Gasiunas et al., 2020).

When finetuned on CRISPR-Cas systems, Evo can coherently generate diverse samples that resemble naturally occuring Cas systems in both sequence and structure. Designing new Cas systems has historically relied on mining sequence databases for homologous proteins, where natural evolution provides functional diversity. Generative modeling with Evo provides an alternative design methodology that can be harnessed across the broad applications of CRISPR technology.

### 2.6. Generative design of transposable biological systems

In addition to molecular complexes, Evo can learn patterns underlying multi-gene systems. An example of minimal replicating systems are mobile genetic elements (MGEs), which are found throughout all domains of life. Their opportunistic spread provides a fundamental force driving sequence variation, new gene function, and even speciation (Chandler et al., 2020). Insertion sequence (IS) elements are compact MGEs that generally encode only the components that are required for transposition. The IS605 group is widely distributed across prokaryotes and consists of three key components: a TnpA transposase that catalyzes peel-and-paste transposition next to an RNA-guided TnpB nuclease and its cognate ωRNA that bias the selfish inheritance of the transposable element (Figure 4A) (Meers et al., 2023; Karvelis et al., 2021; Altae-Tran et al., 2021). The IS605 group belongs to the greater IS200/IS605 family, which includes IS200 group elements that lack the TnpB endonuclease. Improving our understanding of their biological function and generating new MGEs with desired properties could lead to more effective genome engineering tools.

**Figure 4.**
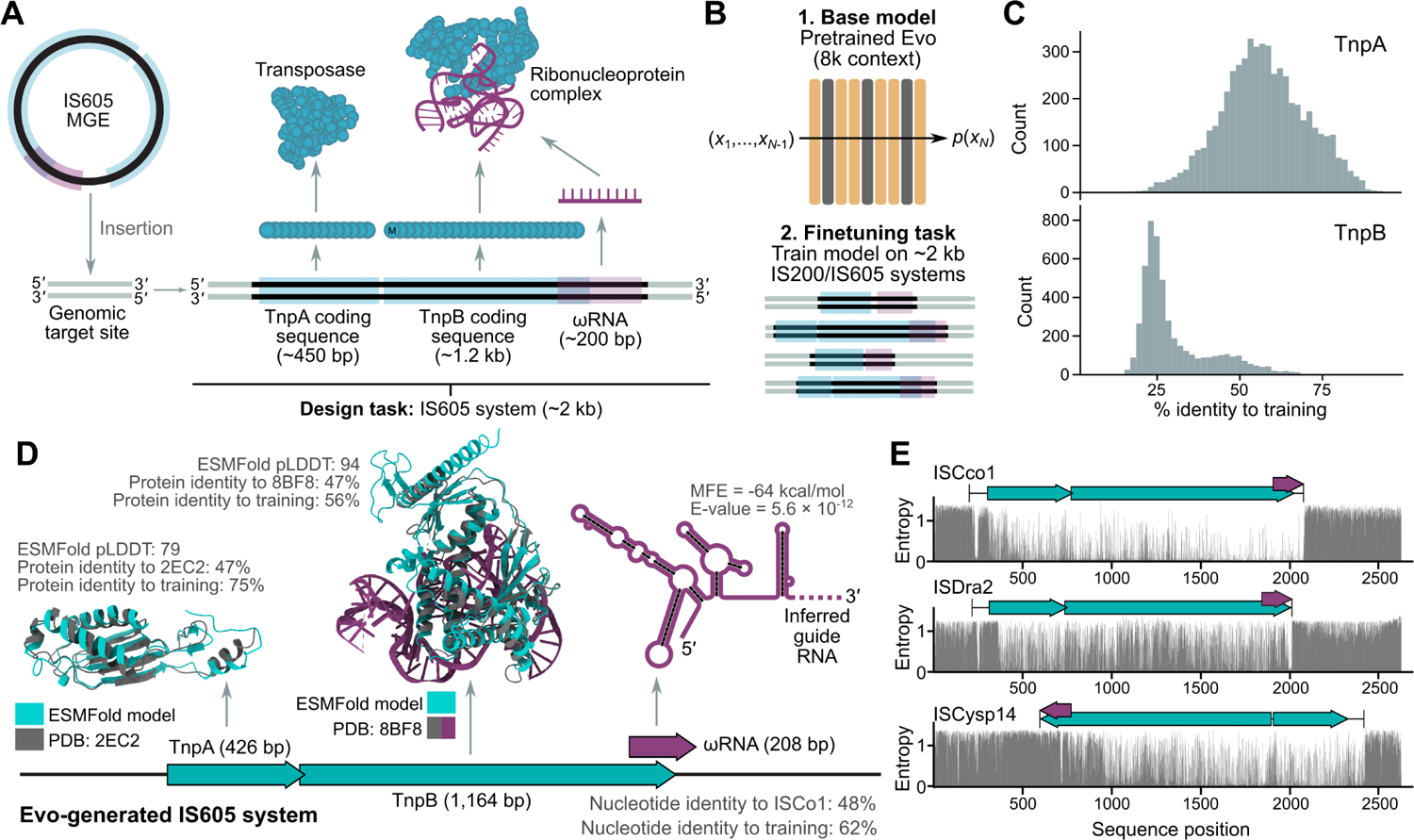
Finetuning on IS200/IS605 sequences enables generative design of transposable biological systems. (**A**) IS605 group systems are a group of MGEs that belong to the IS200/IS605 family and encode a Y1-HuH transposase encoded by the TnpA coding sequence and a TnpB-ωRNA complex that performs DNA cleavage. Our design task is to produce sequences that contain these protein and ncRNA components. (**B**) We finetuned Evo, following its initial 8k pretraining phase, on ∼2 kb-length sequences containing IS200/IS605 systems. (**C**) Histograms representing the distribution of percentage identity of generated loci that contained a predicted TnpA and TnpB coding sequences. The closest matching member of the training set as identified by MMseqs2 was compared with the generated sequence by MAFFT alignment. (**D**) Example of a generated IS605 element. Showing TnpA and TnpB protein structures as predicted by ESMFold (blue) aligned to homologous PDB structures (gray) and the secondary structure of a predicted ωRNA. (**E**) Showing the entropy of the conditional probabilities at each position across three natural IS605 loci. TnpA (short) and TnpB (long) coding sequences are shown in blue, predicted ωRNA boundaries are shown in purple, and the start and end of the complete element is shown with black bars.

We finetuned Evo on 10,720 IS605 elements and 219,867 IS200 elements in their natural sequence context and used the model to generate novel IS200/IS605 elements (Figure 4B) (**Methods**). Focusing on generated sequences that contained both a predicted TnpA and TnpB coding sequence, we successfully detected many sequences that encoded proteins that diverged substantially from the training set, with 22.5% of TnpA proteins being <50% identical to the training set, and 90.1% of TnpB proteins (Figures 4C and **S8**). We found that 87.6% of generated TnpA proteins folded well with ESMFold pLDDT > 70, compared to 25.5% of TnpB proteins, which may be due to the greater abundance of TnpA proteins in the training set. Through annotation and inspection of individual examples, we found that some diverse loci encoded coherent transposase and nuclease proteins that folded well using ESMFold, closely matched experimentally determined structures of homologous proteins, and also contained predicted ωRNA sequences (cmsearch E-value = 5.6 × 10^−12^; Figure 4D).

MGEs are highly abundant and evolve rapidly, making it difficult to systematically identify the precise boundaries of the elements in their natural sequence context (Durrant et al., 2020). Using the finetuned model, we calculated the entropy of the conditional probabilities at each position across natural IS605 loci (Figures 4E and **S8**). Although the model was trained without any explicit labeling of MGE boundaries, the entropy signal indicates that the model is learning a representation of these boundaries, with a sharp and sustained increase in entropy corresponding with the 3^′^ end of the element in particular. Taken together, these results indicate that the finetuned model can generate diverse IS605 systems with coherent protein and RNA sequences, and that the model is learning important features of these elements that could be repurposed for improved functional annotation.

### 2.7. Predicting gene essentiality with long genomic context

Beyond the molecular or systems level, we designed Evo to be capable of analyzing whole genomes. We conducted a second stage of pretraining using the 8k-pretrained Evo model as the base model, training it on sequences of 131k tokens (Figure 5A) with prepended species-level special tokens. This pretraining stage used data from GTDB and a subset of IMG/VR that excludes eukaryotic viruses (Figures 1C and **S1**). See **Methods** for additional details related to context extension. Importantly, Evo maintains single-nucleotide resolution at its 131k context size, which is important because changes involving small numbers of base pairs can still dramatically affect a whole organism’s phenotype. For example, even a single-nucleotide mutation in an essential gene can be incompatible with life if it disrupts that gene’s expression or function. Identifying these essential genes is important for understanding the fundamental biology of an organism and for identifying genes in pathogenic organisms that could be the targets of inhibitory drugs (Rocha and Danchin, 2003).

**Figure 5.**
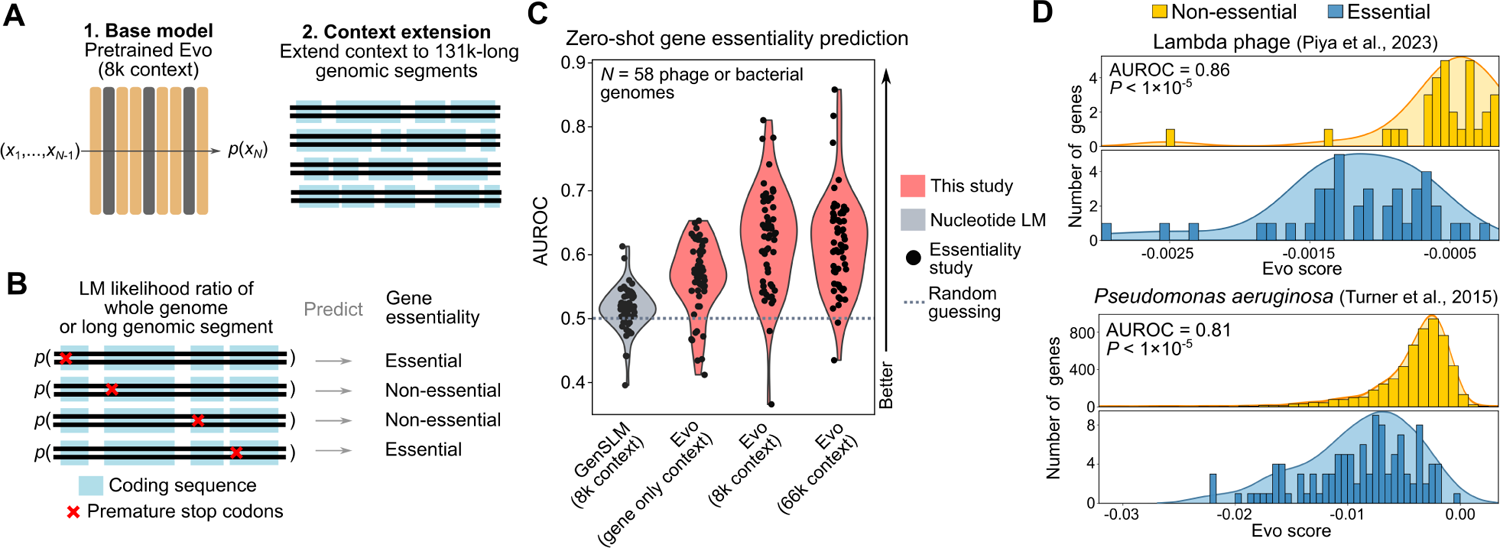
Evo performs zero-shot gene essentiality prediction across diverse bacterial and phage genomes. (**A**) For genome-scale prediction and generation tasks, we first pretrained Evo on sequences with 8k tokens and then extended its context window size in a second pretraining phase to sequences of 131k tokens. (**B**) We performed an in-silico, genome-wide mutagenesis screen in which we introduced premature stop codons at each coding sequence in a genome. We computed the language model (LM) likelihood of the mutated gene sequence plus some amount of additional genomic context (up to 66 kb). We then took the ratio of this likelihood to the likelihood of the unmutated sequence. We tested whether these likelihood ratios would be predictive of gene essentiality. (**C**) Violin and strip plots of the distribution of the strength of gene essentiality prediction across 58 studies (each dot corresponds to a different study), in which each study conducted a genome-wide essentiality screen in a bacterial (*N* = 56) or phage (*N* = 2) species. We measured predictive performance as the AUROC in which the LM likelihood ratio is used to predict a binary label of “essential” or “nonessential.” “Gene only context” indicates that the model is provided with only the gene sequence and no additional flanking genomic context. “8k context” and “66k context” indicate that the LM is provided with the gene sequence and flanking genomic context up to a total of 8k or 66k tokens, respectively. Evo has some predictive performance with gene only context, has vastly improved performance from gene-only to 8k context, and some outlier improvements from 8k to 66 context. (**D**) Histograms representing the distributions of the log of the likelihood ratios (“Evo score”) for the essential genes (blue) and the nonessential genes (yellow) in two genomes: lambda phage (top) and *Pseudomonas aeruginosa* (bottom). These results are based on providing Evo with 66k context.

To this end, we evaluated Evo’s ability to predict gene essentiality solely based on mutations to the genome sequence. We conducted an experiment in which we inserted premature stop codons at the beginning of each coding sequence in a given organism’s genome and measured the effects of these changes on Evo’s likelihood with respect to the likelihood of the wildtype sequence (Figure 5B). When computing the changes to the mutant versus wildtype sequences, we evaluated Evo on the gene sequence alone (“gene only context”), or the gene sequence with flanking context up to a total of 8k tokens (“8k context”) or 66k tokens (“66k context”) (**Methods**). We hypothesized that mutations to essential genes would result in larger, more negative changes in log-likelihood compared to mutations to non-essential genes, allowing us to predict gene essentiality.

On a dataset of 56 whole-genome essentiality studies in bacteria from the DEG database (Zhang, 2004) and two whole-genome essentiality studies in phage from Piya et al. (2023), we observed that the changes in Evo log-likelihood with 66k context are significantly predictive (Bonferroni-corrected permutation-based *P* < 0.05) of gene essentiality in 43 out of 58 genomes. We also observed that providing the model with additional genomic context beyond the gene sequence results in a substantial improvement in performance, especially from gene only context to 8k context. From 8k to 66k context, the average predictive performance is essentially equivalent, but the range does increase due to improvement in outlier examples (Figures 5C and **S9A**). With 8k context, the model most likely has access to enough of the genome to improve its prediction of mutational effects on organism function, whereas 66k context provides new, helpful information in only some cases. For a few genomes, the zero-shot performance with 66k context is notably strong, with an AUROC of 0.86 on lambda phage essentiality data (Piya et al., 2023) and an AUROC of 0.81 on *Pseudomonas aeruginosa* essentiality data (Turner et al., 2015) (Figure 5D).

Evo is also able to predict essentiality when using different in-silico mutagenesis strategies, such as varying the number of stop codons inserted, or deleting the gene sequence entirely (**Figure S9B**; **Methods**), though we did not attempt an exhaustive search of the best prompting strategy for this task. GenSLM, a codon language model that had mild predictive performance of mutational effects on single-gene protein function (Figure 2B), could not perform this zero-shot prediction task (Figure 5C). We also observed that a gene’s position in the genome is not predictive of essentiality, indicating that trivial positional positional biases do not contribute to prediction performance (**Figure S9B**). Together, these results demonstrate that Evo can predict mutational effects at a whole-organism level across many bacterial and phage species, without any explicit genome annotations, task-specific training data, or functional labels. In contrast to protein or codon language models, Evo enables an understanding of gene function within a broader genomic context.

### 2.8. Generating DNA sequences at genome scale

Given Evo’s generative capabilities, we were interested in testing its generation quality at long sequence lengths without additional finetuning. By doing so, we can better understand the patterns and the level of detail learned by the model, which helps us determine the model’s capabilities and limitations. We used Evo to sample twenty sequences each containing ∼650 kb, representing about five times the model’s context length of 131 kb. For comparison, the smallest “minimal” bacterial genomes are about 580 kb in length (Blanchard and Bébéar, 2011). We prompted the model to generate bacterial genomes using the species-level tokens in the training dataset (Figure 6A). To analyze how well these generations recapitulate natural genomes, we used CheckM (Parks et al., 2015), a tool originally developed to assess the quality of bacterial DNA sequenced from nature. CheckM calculates statistics such as the density of coding sequences in the genome and the presence of key marker genes that are found in nearly all prokaryotes, which we used to determine how well our generated sequences mirror key characteristics of natural genomes.

**Figure 6.**
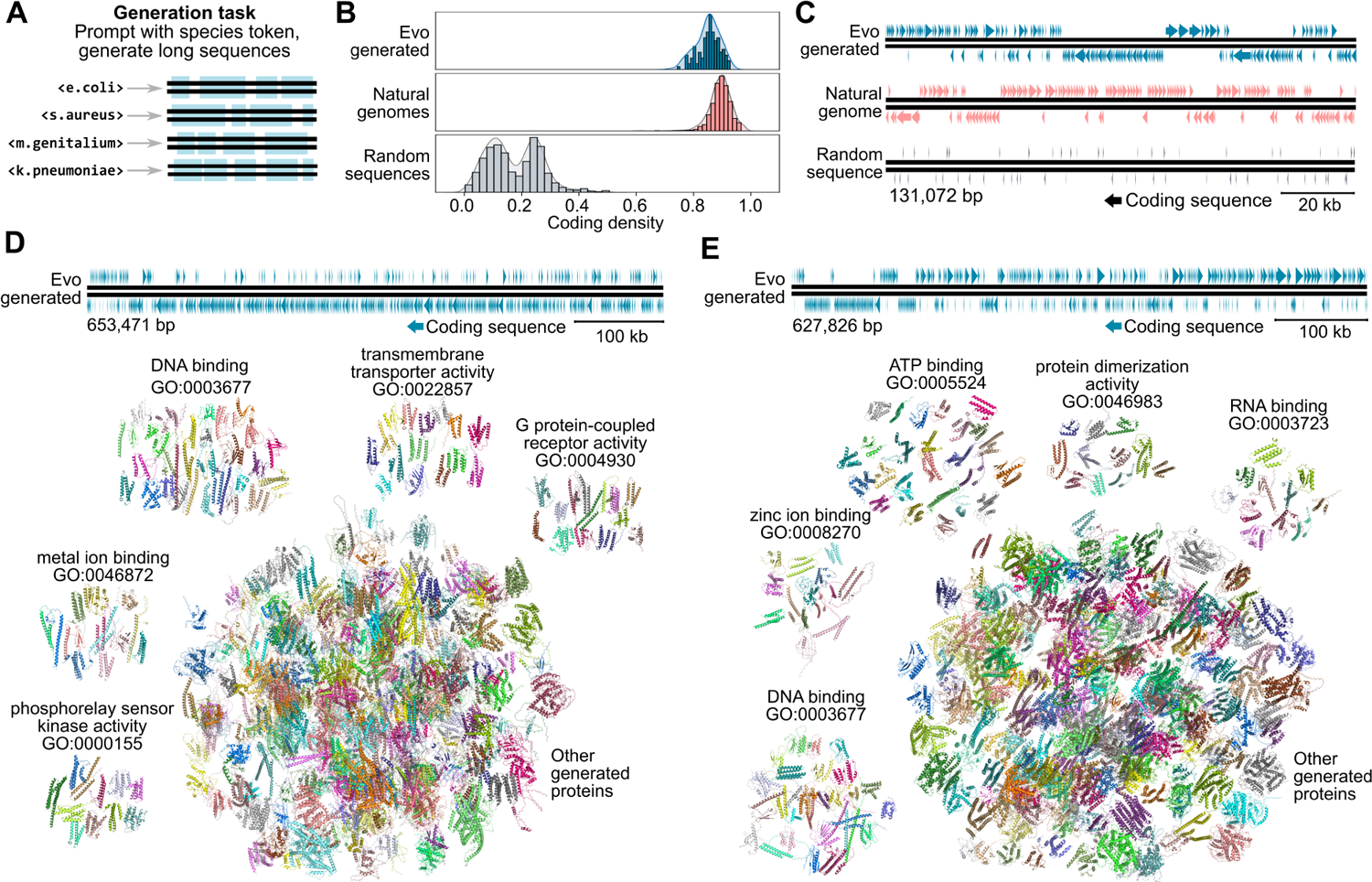
Evo generates genome-scale sequences with dense coding architecture. (**A**) We prompted Evo with species-level tokens used during the second pretraining stage. We use bacterial species prompts and generate sequences of ∼650 kb in length. (**B**) Histograms depicting the distribution of coding density scores among 131 kb crops of sequences generated by Evo (“Evo generated”), sequences from natural bacteria (“natural genomes”), or sequences in which the four base pairs were sampled uniformly at random (“random sequences”). (**C**) Arrow plots depicting the organization of coding sequences on an example 131 kb sequence generated by Evo, derived from a natural genome, or sampled randomly. Coding sequences are depicted as arrows in which the horizontal length of the arrow corresponds to the genomic interval and the direction of the arrow indicates the strand. The top and bottom rows of arrows indicate the 5^′^-to-3^′^ and 3^′^-to-5^′^ strands, respectively, and the Evo-generated sequence was designated as the 5^′^-to-3^′^ strand. Both Evo-generated and natural genomes exhibit operon-like structure in which clusters of co-located genes are on the same strand. (**D**, **E**) Example generated sequences are represented as arrow plots as in (**C**). Below these arrow plots are ESMFold structure predictions of all protein coding sequences from 100 through 1024 amino acids in length as identified by Prodigal. Structure predictions are aligned to natural proteins, which are then mapped to associated GO molecular function terms (**Methods**). The largest GO categories are displayed as clusters alongside a large cluster containing all other proteins.

Notably, the coding density of sequences generated by Evo is nearly as high on average as the density of coding sequences found in natural genomes, and is substantially higher than the coding density of random sequences (Figure 6B). Importantly, when visualized, both natural and generated sequences display similar patterns of coding organization (Figure 6C), with sequences in close proximity typically found with the same strand orientation. In bacteria, these closely linked groups of coding sequences typically correspond to functionally tied gene clusters or operons. When using ESMFold to obtain protein structure predictions corresponding to these coding sequences, almost all showed some predicted secondary structure and globular folds (Figures 6D, **6E**, and **S10**). Some proteins also showed structural similarity to natural proteins involved in known molecular functions as annotated by Gene Ontology (GO) (Figures 6D and **6E**). However, many of these structure predictions are of low confidence and have limited structural matches to any entry in a representative database of naturally occurring proteins (**Figure S10**). The generated sequences also do not contain many highly conserved marker genes that typically indicate complete genomes. Across all of our generated sequences representing ∼13 megabases, Evo sampled 18 tRNA sequences (compared to 35 tRNAs in the ∼580 kb genome of *M. genitalium*) and no rRNAs, as detected by the programs tRNAscan-SE (Chan and Lowe, 2019) and barrnap (Seemann, 2018), respectively.

These results suggest that Evo can generate genome sequences containing plausible high-level genomic organization at an unprecedented scale without extensive prompt engineering or finetuning. These samples represent a “blurry image” of a genome that contains key characteristics but lacks the finer-grained details typical of natural genomes. This is consistent with findings involving generative models in other domains, such as natural language or image generation. For example, directly sampling from a large natural language model typically produces sequences that are grammatically correct yet locally biased toward simpler sentence constructions and that are globally incoherent, especially at long lengths. Promisingly, in these domains, algorithmic techniques have emerged to improve the quality of generations compared to sampling from the pretrained model alone (Wei et al., 2022; Ouyang et al., 2022; Rafailov et al., 2024). The baseline generation quality observed for pretrained Evo suggests that Evo is also amenable to these techniques.

## 3. Discussion

Evo is a genomic foundation model trained on hundreds of billions of DNA tokens across the evolutionary diversity of prokaryotic life, capable of prediction and generation tasks at the scale of individual molecules, molecular complexes, biological systems, and even whole genomes. Based on a state-of-the art hybrid model architecture, StripedHyena, Evo enables single-nucleotide resolution language modeling at a context length of 131k. We conducted the first scaling laws analysis of DNA pretraining across several architectures, where we observed StripedHyena outperforming several baseline architectures, including the Transformer architecture, at each level of scale. Evo accurately performed zero-shot prediction across diverse fitness or expression prediction tasks on proteins, ncRNAs, or regulatory DNA that matches or outperforms specialized models, while also understanding which genes are essential to organismal fitness. Evo is also a generative model, which we leverage to sample CRISPR-Cas proteins and their noncoding guide RNAs, multi-gene transposable systems, and ∼650 kb sequences that recapitulate the coding organization of real genomes. We make open-source code and models for Evo publicly available at https://github.com/evo-design/evo.

A model capable of genome-scale design holds great potential to advance therapeutic discovery, sustainability, and our understanding of fundamental biology, but simultaneously raises biosafety and ethical considerations. The Global Alliance for Genomics and Health (GA4GH) (Rehm et al., 2021) has developed principles for the oversight of genetic engineering technologies and could provide a robust foundation for transparency, accountability, and shared responsibility. Such a framework is essential to foster international cooperation that benefits all humanity. A proactive discussion involving the scientific community, security experts, and policymakers is imperative to prevent misuse and to promote effective strategies for mitigating existing and emerging threats. Furthermore, investment in education and capacity building, especially in under-resourced communities, will sustainably democratize access and use of tools like Evo. We open source the model to promote transparency and begin a dialogue with the broader scientific community and shareholders. We also apply the precaution of excluding eukaryotic viruses from our pretraining dataset. We include an extended discussion on ethical considerations in a supplementary **Safety and ethics discussion** in which we assessed the risks of tools like Evo, including their potential to be misused, contribute to social and health inequity, or disrupt the natural environment. Clear, comprehensive guidelines that delineate ethical practices for the field are required for the responsible development and use of genome-scale language models.

Despite the remarkable capabilities of this first-generation DNA foundation model, a number of technical limitations and challenges remain. We pretrained Evo on a dataset of 300B prokaryotic tokens which represents a miniscule portion of petabytes of publicly available genomic data. Because our model is trained only on prokaryotic data, our ability to predict functional effects of mutations on human protein fitness is limited. Natural language models often struggle to maintain coherent and diverse generation over long sequences, and Evo can demonstrate similar properties. For example, we observed that more novel CRISPR-Cas sequences were sampled at relatively low frequency and prompting on special tokens had moderate controllability, occasionally generating a Cas9 protein when prompted with a Cas12 token. At the genome-scale, Evo generates hundreds of kilobases that demonstrate a high-level understanding of genome organization, but struggles to include key marker genes such as full tRNA-encoding repertoires. These limitations mirror the constraints of natural language models, which have been improved over time with increased scale, labeled data, prompt engineering, and alignment with human preferences (Kaplan et al., 2020; Ouyang et al., 2022; Wei et al., 2022; Kojima et al., 2022; Rafailov et al., 2024). We expect a similar trajectory for models of DNA.

DNA modeling at this scale and resolution lays the groundwork for a host of research directions. We expect that Evo will benefit from additional scale, longer context length, and more diverse pretraining data. Given the success of language-model-guided directed evolution of proteins (Hie et al., 2024), genomic language models may also help guide the directed evolution of multi-gene biological systems. Similarly, the co-evolutionary information contained in these models could improve molecular structure prediction in a multi-gene context (Jumper et al., 2021; Lin et al., 2023). Properties of systems biology may emerge as these models improve, such as fitness effects of combinatorial gene interactions or the prediction of functional operon linkages. With better conditioning or prompt engineering, Evo could form the basis of a next-generation sequence search algorithm by enabling metagenomic mining at a relational or a semantic level rather than extracting literal sequences from existing organisms. Beyond prokaryotes, the incorporation of eukaryotic genomes into Evo will need to consider the far higher complexity of these genomes and require substantial resource investment in engineering, compute, and safety-related model alignment. Combined with advances in large-scale genome modification (Durrant et al., 2024), Evo helps expand the scope of biological engineering and design to the scale of whole genomes.

## 4. Code and data availability

Code and models related to this study are publicly available at https://github.com/evo-design/evo. We used the following datasets for pretraining:

- Bacterial and archaeal genomes from the Genome Taxonomy Database (GTDB) v214.1 (Parks et al., 2015).
- Curated prokaryotic viruses from the IMG/VR v4 database (Camargo et al., 2023).
- Plasmid sequences from the IMG/PR database (Camargo et al., 2024).

In addition to the above datasets, we also used portions of the following datasets for finetuning:

- NCBI RefSeq (O’Leary et al., 2016).
- UHGG (Almeida et al., 2021).
- JGI IMG (Chen et al., 2021).
- The Gut Phage Database (Camarillo-Guerrero et al., 2021).
- The Human Gastrointestinal Bacteria Genome Collection (Forster et al., 2019).
- MGnify (Mitchell et al., 2020).
- Youngblut et al. (2020) animal gut metagenomes.
- MGRAST (Meyer et al., 2008).
- Tara Oceans samples (Sunagawa et al., 2015).

Additional details on these datasets are provided in **Methods**.

## Acknowledgements

We thank Elijah Chanakira, Dave Driggers, Richard Dugan, Helmut Fritz, Marco Iskender, Adeesh Jain, Mike LaPan, Sean Marrs, Sigalit Perelson, Randy Rizun, Jason Rojas, and Delaney Ugelstad for assistance with computational infrastructure. We thank Samuel Sternberg and Chance Meers for providing covariance models to identify diverse ωRNAs. We thank Jessica Adkins, Joana Carvalho, Dan Fu, Jared Dunnmon, Yunha Hwang, Julia Kazaks, Gautam Machiraju, April Pawluk, Christina Theodoris, Ben Viggiano, and Alden Woodrow for helpful discussions and assistance with manuscript preparation. P.D.H. acknowledges funding support from Arc Institute, Rainwater Foundation, Curci Foundation, Rose Hill Innovators Program, V. and N. Khosla, S. Altman, and anonymous gifts to the Hsu Lab. B.L.H. acknowledges funding support from Arc Institute, Varun Gupta, and Raymond Tonsing.

## 5. Author Contributions

E.N., P.D.H., and B.L.H conceived the project. P.D.H. and B.L.H. supervised the project. E.N., M.P., and A.W.T. designed the model architecture. M.G.D. and B.L.H. curated and processed the pretraining and finetuning datasets. M.P. implemented the optimized training and generation infrastructure. E.N., A.W.T., and B.L.H. contributed to the optimized training and generation infrastructure. E.N., M.P., and A.W.T. implemented and carried out the scaling laws analysis. E.N., M.P., A.W.T., and B.L.H. evaluated the pretrained model. E.N. and B.L.H. conducted model finetuning. B.K. and P.D.H. sampled or analyzed CRISPR-Cas generations. M.G.D. and B.L.H. sampled or analyzed the IS200/IS605 generations. B.L.H. conducted the gene essentiality analysis. M.P. and B.L.H. conducted genome-scale sampling and analysis. M.P., A.W.T., and B.L.H. implemented the public Evo codebase. M.Y.N., A.L., and T.H-B. conducted the ethics and safety investigation and discussion. E.N., M.P., M.G.D., P.D.H., and B.L.H wrote the first draft of the manuscript. All authors wrote the final draft of the manuscript.

## Supplementary Material

### A. Safety and ethics discussion

The introduction of powerful generative genomic foundation models such as Evo enables the rapid deciphering of complex genetic information, which can be used for genetic engineering and therapeutic development. Evo is the first of its kind to predict and generate DNA sequences at the whole-genome scale with single-nucleotide resolution, albeit only for prokaryotes in this version. As future capability increases are likely achievable with the class of large-scale DNA models enabled by Evo, we provide an extended ethical discussion on potential risks and precautionary measures. While Evo is limited in its current form, the molecular design, synthesis, manipulation, and dissemination of new synthetic genetic materials could pose concerns to individuals, society, and the environment. Through a responsible AI lens (Badal et al., 2023), we forecast three salient ethical implications and identify mediating solutions.

### A.1. Safety and ethical implications

#### A.1.1. Whole-genome foundation models have the potential for misuse

There are concerns that the dual-use (or misuse) of genomic foundation models by malevolent actors could pose a threat to biosafety and biosecurity (Baker and Church, 2024). Tools like Evo serve to enhance queries of the existing genomic knowledge base and identify genetic regions of interest for editing or experimentation. The ability to discern fitness associated with certain sequences can assist in the discovery of novel biomarkers or therapeutic targets, but can also catalyze the development of harmful synthetic microorganisms that more easily bypass the body’s natural defenses, are resistant to current treatments, or cause more severe disease. Fortunately, even with optimal synthetic genomic designs, the ability to create viable organisms is limited by high barriers to entry, including a substantial amount of technical resources and expertise needed to carry out genome synthesis and expression, which is further compounded by the unpredictability of biological mechanisms. Nevertheless, as genetic engineering tools become more readily available, guardrails (for example, access controls, usage audits) should be agreed upon by shareholders to limit unfettered queries for harmful genetic sequences. Clear definitions of what constitutes “dual-misuse” are also needed to draw the line for researchers, policy makers, and other shareholders.

#### A.1.2. Whole-genome foundation models could contribute to social and health inequity

Given the high barriers to entry, access and capability inequality with tools such as Evo can lead to inadvertent societal harms. Evo is open source to promote transparency and reproducible research. However, those who can most effectively use, and hence benefit the most from, the tool are entities with coordinated biotechnical resources and expertise, such as biotechnology and pharmaceutical corporations. These companies may accelerate research in a direction that prioritizes returns-on-investment over the global disease burden or health equity (Morin et al., 2023). Along the same line, wealthier nations or more well-funded institutions also stand to better leverage Evo to accelerate their research agendas, further widening the gap between high- and low-resource settings.

The use of generative tools in biology also raises complex intellectual property concerns. Biological foundation models such as Evo may enable an organization to bypass current intellectual property limitations on biological therapeutics or other materials. In some cases, this may lead to a monopolization of treatments for certain conditions. Such an entity could then use these rights to set prohibitively high prices and make treatments inaccessible to most patients (for example, those in low-income countries), thus further exacerbating health disparities (Peek, 2021). In other cases, bypassing intellectual property protections could discourage further investment into therapeutic innovation. Overall, we argue that an entity that uses and benefits from open-source tools such as Evo has a duty to return value to the public and contribute to social and health equity. Intellectual property law should also evolve as generative models increasingly automate the biological discovery and design process.

#### A.1.3. Whole-genome foundation models could contribute to disruptions to the natural environment

Although Evo does not directly manipulate any genetic material, it may enhance the efficiency of genetic engineering projects. There are concerns with how the capabilities of genetic engineering technologies may disrupt the environment and cause ecological uncertainty (for example, the release of altered organisms), leading to a loss of biodiversity or the emergence of new, potentially harmful species (Macfarlane et al., 2022). Although the ecological impacts of training whole-genome foundation models remain unknown, more immediately, it is also important to consider the carbon footprint associated with increasing infrastructure and computational demands (Nature Computational Science, 2023). The capabilities of tools such as Evo, alongside other technologies for genome editing and ecological engineering, add to complex debates about the extent to which science should intervene in evolution. As we push the boundaries of scientific capabilities with tools such as Evo, it becomes imperative to reflect on the interactions and boundaries between our inventions and natural evolutionary processes, aiming to preserve ecological balance, maintain environmental sustainability, and uphold ethical standards.

### A.2. The path forward

The path forward for the responsible use and development of tools like Evo is anchored in the establishment of clear, comprehensive guidelines that delineate ethical practices. These guidelines serve as a responsible AI framework, ensuring that all shareholders—researchers, developers, and users—have a common understanding of the safety and ethical dimensions inherent in genetic engineering. Coupled with robust oversight mechanisms, this approach aims to monitor and manage the application of Evo to prevent misuse and ensure its alignment with ethical standards. Furthermore, promoting transparency regarding the use of these technologies and fostering open dialogue among all parties will enhance trust and collaboration within the scientific community and beyond.

To address disparities in access and capabilities, particularly in low-income countries, the strategy includes forging community partnerships and international collaborations. By offering targeted training and support, these partnerships can democratize access to advanced tools like Evo, enabling a broader spectrum of scientists and researchers to contribute to and benefit from genetic engineering innovations. At the policy level, investing in education and capacity building emerges as a pivotal element, equipping the next generation of scientists with the ethical acumen and technical skills to navigate the complexities of genetic research responsibly.

Central to sustaining ethical innovation is the creation of a dynamic feedback loop that engages all shareholders in a continuous dialogue. By setting up mechanisms to collect and integrate feedback from those involved in or impacted by Evo’s applications, the process ensures that guidelines, policies, and practices are regularly refined in response to evolving ethical challenges and societal expectations. Collaborating with organizations such as the Global Alliance for Genomics and Health (GA4GH) (Rehm et al., 2021) to develop and update genetic engineering guidelines further solidifies this commitment to ethical excellence. This multifaceted approach not only addresses immediate concerns but also lays the groundwork for a future where genetic engineering advances in harmony with ethical principles and societal values.

## B. Methods

### B.1. StripedHyena architecture

Evo is based on StripedHyena (Poli et al., 2023a), a state-of-the-art hybrid model architecture for sequence modeling. Evo comprises 32 blocks at a model width of 4096 dimensions. Each block contains a sequence mixing layer, tasked with processing information along the sequence dimension, and a channel mixing layer, focused on processing information along the model width dimension. In the sequence mixing layers, Evo employs 29 hyena layers, interleaved with 3 rotary (Su et al., 2024) self-attention layers at equal intervals. We parametrize convolutions in hyena operators using the modal canonical form described in (Massaroli et al., 2024). For the channel mixing layers, Evo employs gated linear units (Dauphin et al., 2017; Shazeer, 2020). Evo further normalizes the inputs to each layer using root mean square layer normalization (Zhang and Sennrich, 2019).

#### Hyena layers

Hyena (Poli et al., 2023a) is a sequence mixer implementing an input-dependent (data-controlled) operator via a composition of short convolutions, long convolutions and gating (Figure 1B). Hyena belongs to the class of deep signal processing primitives (Poli et al., 2023a; Fu et al., 2024; Massaroli et al., 2024), designed for efficient, input-dependent computation in large-scale sequence models. Input-dependence enables an architecture built with deep signal processing layers to adapt computation based on the input, enabling in-context learning (Arora et al., 2023; Bhattamishra et al., 2023). These layers rely on structured operators compatible with fast multiplication algorithms and can thus be evaluated in subquadratic time using, e.g., Fast Fourier Transforms for convolutions. The operators are parametrized *implicitly*, e.g., learning a map from positional embeddings, or the input, to the parameters of the operator. Typical choices of implicit parametrizations are linear projections, hypernetworks (Romero et al., 2021; Poli et al., 2023a) or linear state-space models in modal or companion form (Gupta et al., 2022; Gu et al., 2022; Massaroli et al., 2024; Orvieto et al., 2023; Zhang et al., 2023). The blueprint of a hyena operator forward pass is summarized below.

**Algorithm 1.**
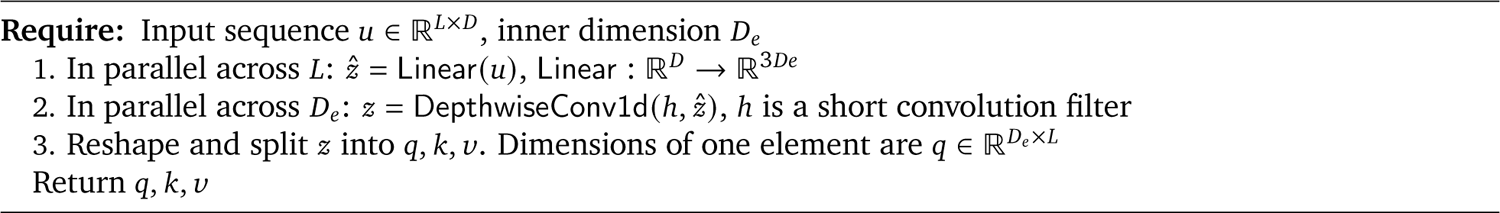
ConvProjection.

**Algorithm 3.**
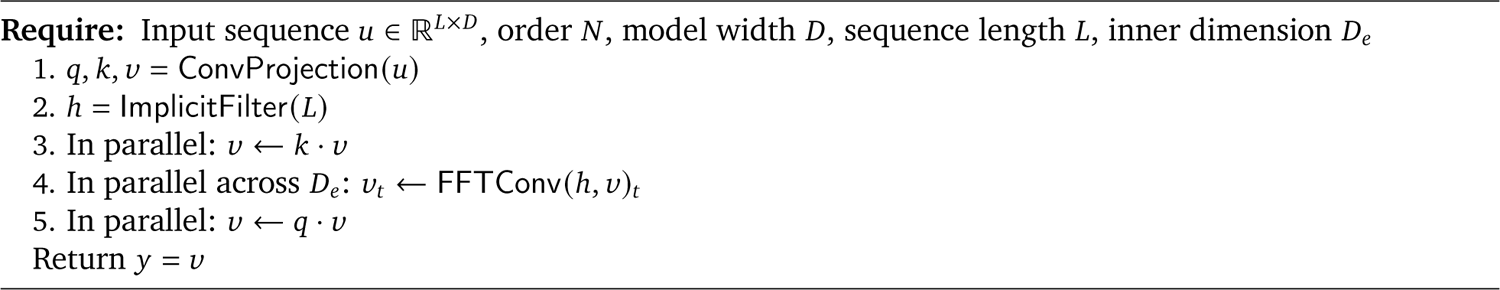
Forward pass.

#### Self-attention layers

Self-attention is the core sequence mixing operator of Transformer models. Self-attention constructs the output sequence as a weighted combination of the input elements, where the weights themselves are input-dependent. Given an input sequence, the forward pass of an (unnormalized) self-attention layer is:

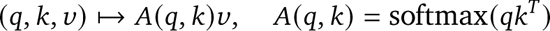

where queries *q* ∈ ℝ^*L*×*D*^ and keys *k* ∈ ℝ^*L*×*D*^ and values *v* ∈ ℝ^*L*×*D*^ are obtained through a linear transformation of the input e.g., *v* = *uW*_*v*_. The softmax is applied to rows of *A*. The query, key, value terminology is borrowed from databases, where keys are used to index stored values. Conceptually, the values of the attention matrix *A*(*q*, *k*) measure the similarity beween queries and keys akin to matching queries to keys in a database.

#### Positional embeddings

By itself, the self-attention operator does not have any notion of the different positions of the input embeddings in an input sequence. For this reason, it is generally supplemented with a positional encoding mechanism. The attention layers of StripedHyena utilize a rotary position embedding mechanism (RoPE) to model relative positional information (Su et al., 2024). Position information is encoded by rotating the query and key token vectors of the attention operator. Specifically, RoPE implements a rotation to queries and keys, with the rotation magnitude defined as a function of their relative position in the sequence.

To extend the context window length from 8k to 131k during our second pretraining stage, we apply linear position interpolation to extend the rotary position embedding applied in the first pretraining stage at 8k sequence length (for details, see Chen et al. (2023)). Interpolating enables the model to continue leveraging its learned representations when applied to longer sequences than it was originally trained on. We also tested other position interpolation methods but found that they performed slightly worse than linear interpolation on our data.

#### Tokenization

In language modeling, tokens describe the smallest unit of semantic information that is used by a model to process language. For example, tokens can indicate individual words of a vocabulary or even lower-level semantic information such as individual characters. Tokenization describes the process of mapping these semantic language units, such as words or characters, to unique integer values, each indicating an entry in a lookup table. These integer values are mapped by embedding layers to vectors, which are then processed by the model in an end-to-end fashion. Evo tokenizes DNA sequences at single-nucleotide resolution, using the UTF-8 encoding implemented in Python. During pretraining, Evo uses an effective vocabulary of four tokens, one per base, from a total vocabulary of 512 characters. We use the additional characters to enable prompting with special tokens during generation with finetuned models.

### B.2. OpenGenome datasets

The OpenGenome pretraining dataset (S3 for summary statistics) was compiled from three different sources:

1. Bacterial and archaeal genomes from the Genome Taxonomy Database (GTDB) v214.1 (Parks et al., 2015),
2. curated prokaryotic viruses from the IMG/VR v4 database (Camargo et al., 2023), and
3. plasmid sequences from the IMG/PR database (Camargo et al., 2024). For GTDB, representative genomes for each species were retained to reduce data redundancy.

For IMG/PR, only one representative per plasmid taxonomic unit (PTU) was kept. For IMG/VR, sequences were retained only if they were labeled as “High-confidence” according to the database metadata, and only one representative per viral operational taxonomic unit (vOTU) was kept. These sequences were further curated to remove potential eukaryotic viruses by keeping only sequences whose assigned taxonomic classification was found within a prokaryotic host at least twice. Next, the remaining taxonomic classifications were inspected and further filtered to exclude all viruses assigned to any of 19 families (Adenoviridae, Caliciviridae, Coronaviridae, Filoviridae, Flaviviridae, Hantaviridae, Hepadnaviridae, Herpesviridae, Orthomyxoviridae, Papillomaviridae, Paramyxoviridae, Picornaviridae, Poxviridae, Reoviridae, Retroviridae, Rhabdoviridae, Circoviridae, Geminiviridae, Picobirnaviridae) or 12 orders (Amarillovirales, Durnavirales, Geplafuvirales, Herpesvirales, Lefavirales, Ortervirales, Orthopolintovirales, Piccovirales, Picornavirales, Priklausovirales, Cirlivirales, Mulpavirales). Next, viruses with poor taxonomic specificity were excluded, including those with no assigned realm at all, and those only assigned up to the level of r:Riboviria, r:Monodnaviria, k:Heunggongvirae, k:Bamfordvirae, p:Preplasmiviricota, p:Cressdnaviricota, p:Pisuviricota, or c:Tectiliviricetes.

The CRISPR/Cas and IS200/IS605 fine-tuning datasets were compiled from a previously described custom database gathered from multiple sources (Wei et al., 2023). Briefly, this custom database includes genomic and metagenomic sequence data from NCBI RefSeq O’Leary et al. (2016), UHGG (Almeida et al., 2021), JGI IMG (Chen et al., 2021), the Gut Phage Database (Camarillo-Guerrero et al., 2021), the Human Gastrointestinal Bacteria Genome Collection (Forster et al., 2019), MGnify (Mitchell et al., 2020), Youngblut et al. (2020) animal gut metagenomes, MGRAST (Meyer et al., 2008), and Tara Oceans samples (Sunagawa et al., 2015).

To compile the CRISPR/Cas genomic loci, this custom database was searched using profile HMM models and the HMMER software package to identify Cas9, Cas12, and Cas13 sequences (Finn et al., 2011). Several pHMMs were collected from the CRISPRCasTyper annotation tool (Russel et al., 2020), and a recent computational survey of TnpB and Cas12 (Altae-Tran et al., 2023). Custom Cas13 pHMMs that were previously generated by our group were also used (Wei et al., 2023). These models were searched against our large custom database using hmmsearch and the parameter “-Z 1000000”. All hits that met E < 1 × 10^−6^ with at least one pHMM were kept. Only hits that were at least 300 aa long and covered over 80% of the pHMM were kept.

For all hits to a given pHMM, only proteins that were within the middle 99% of the size distribution were kept. Corresponding genetic loci were extracted from the database, including 8,192 nucleotides of flanking sequence on both the 5^′^ and 3^′^ ends of the Cas effector CDS. The tool minced was used to identify CRISPR arrays in the flanking sequences using the parameters “-minRL 18 - maxRL 50 -minSL 18 -maxRL 50” (?). Only loci with both a predicted Cas effector and a CRISPR array were retained. The final CRISPR/Cas loci were extracted by first identifying the subsequence that covered both the Cas effector and the CRISPR array, and then including additional flanking nucleotides on both sides up until 8,192 were retained for fine-tuning purposes. Only 1 locus per 90% identity Cas cluster was retained, clustered using the MMseqs2 command “easy-cluster ––cluster-reassign ––cluster-mode 0 ––cov-mode 0 -c 0.7 ––min-seq-id 0.9” (Steinegger and Söding, 2017).

To compile the IS200/IS605 loci, this custom database was searched using a Pfam Y1 HUH Transposase pHMM model (Pfam ID: PF01797). This pHMM identifies IS200/IS605 TnpA proteins. All matches meeting E-value < 1 × 10^−6^ that covered at least 80% of the pHMM and were less than 400 aa were kept. 8,196 nt of CDS-flanking sequence was then extracted for each hit. Loci that also contained TnpB coding sequences were identified using previously compiled pHMMs (Altae-Tran et al., 2023), and a custom pHMM compiled using jackhmmer and the ISDra2 TnpB as an initial query against the MGnify protein database, followed by a MAFFT alignment of hits and pHMM construction with HMMER (Finn et al., 2011; Mitchell et al., 2020; Katoh et al., 2002). Hits that were between 250 and 650 aa in length were retained, and only loci where the distance between the beginning and end of the TnpA and TnpB sequences was less than 2500 nt were retained. For TnpA-only loci, up to 300 nt of flanking sequence were added to either side of the CDS. For TnpA+TnpB loci, up to 300 nt were added to the TnpA side of the IS200/IS605 element, while 600 nt were added to the TnpB side (to account for the presence of an ωRNA). Only 1 locus per 90% identity TnpA cluster was retained.

### B.3. Training procedure

We pretrain Evo in two stages, first with a context size of 8k tokens, followed by a second stage where we increase the context size to 131k tokens. Multi-stage sequence length pretraining has been shown to reduce the overall number of compute hours required to train long context models (Xiong et al., 2023). In total, we trained Evo in stage 1 on 64 Nvidia H100 GPUs and on 128 Nvidia A100 GPUs in stage 2. In total, Evo was trained on approximately 340B tokens, using approximately 2 × 10^22^ FLOPS. For specific generation tasks, we further finetuned Evo, as described in the following sections. We also report long context perplexity scaling of Evo 131k in Figure S2. Additional details on training are provided in Table S2.

#### Dataloading

We use sequence packing to generate training samples, where multiple DNA sequences are appended until the context length (8k or 131k) is reached. Individual DNA sequences are separated by end-of-sequence (EOS) tokens. Depending on the dataset or task, we additionally prepend a special class (CLS) token to condition the model, for example, to steer its generations through prompting.

#### Hyperparameter tuning and direct model comparisons

Before training Evo, we carried out hyperparameter tuning on partially trained 7B Transformer++ (see B.4) models and compared to similarly sized Hyena and StripedHyena models. In particular, we swept batch size, learning rate and other architectural details. Even when controlling for training iterations instead of compute (FLOPS), Transformer++ performance is substantially worse than StripedHyena (see S4). Out of all the baselines, we find that StripedHyena achieves the overall lowest perplexity at the 7B scale, consistent with the scaling rates presented in Figure 1G.

### B.4. Scaling laws

We compare different classes of architectures via a compute-optimal protocol, aimed at evaluating results on the *compute-optimal frontier*. Compute-optimal analysis studies the best performance of a pretraining run given a compute budget, typically indicated in floating point operations (FLOPs), and achieved by optimally allocating portions of the compute budget to model size and dataset size. Architecture types differ in compute efficiency, as well as how they allocate this compute budget.

We started by tuning hyperparameters such as learning rate and batch size for Transformer++ with a grid search, then used the same values for all architectures except in settings where numerical instability was observed. To address instability, we lowered the learning rate gradually and repeated the experiment until convergence. In all experiments, we trained models with 8,192 tokens in context length. For each compute budget defined by a total FLOP count, we varied the model sizes (6 million to 1 billion parameters) and the number of tokens trained. To measure model performance, we use the perplexity metric, which indicates how well an autoregressive model performs at predicting the next token of a sequence and is highly correlated with performance on downstream tasks. A lower perplexity value indicates better performance.

#### Scaling laws procedure

We provide a summary of the steps involved in our scaling laws analysis. Quantifying scaling rates allows us to predict performance as model size, dataset size, and compute grow.

1. Define a set of compute budgets to study. We use 8 × 10^18^, 2 × 10^19^, 4 × 10^19^ and 8 × 10^19^ FLOPS.
2. Calculate the FLOPS (floating point operations) required to process a fixed input size for the model architecture of interest (i.e. the “cost” of using the model).
3. Identify the model’s compute-optimal allocation for each compute budget:

(a) Select a wide range of possible model sizes, and calculate for each model size the corresponding number of tokens that need to be processed to reach the compute budget. Other hyperparameters are chosen according to Table S1. We generally observe minor changes to model topology (depth, width) to only minimally affect perplexity, aligning our results with the findings presented by Kaplan et al. (2020) for Transformers.
(b) Train a model of each size and record its performance (e.g., in terms of perplexity).
(c) Identify the optimal compute allocation: Following prior analysis, we fit a second-order polynomial as a function from (log) model size to perplexity, and extract obtained the compute-optimal point as its minimum. The compute-optimal point identifies the optimal allocation of model size and training tokens at the given compute budget.

After deriving the compute-optimal scaling rates (Figure 1G), we compare architectures and compute optimal allocation of tokens and model size (Figure S5). In Figure S3, we also show rates for compute-suboptimal model sizes by architecture. In particular, we quantify the effect on perplexity scaling caused by a suboptimal allocation of compute budget to model or dataset size (e.g., training a smaller model for more tokens). We estimate the compute-optimal model size for each compute budget, then reduce it by a percentage (the *offset*). The corresponding perplexity is obtained via the IsoFLOP curves (Figure 1F). Transformer++ perplexity scaling rapidly degrades outside the compute-optimal frontier, in contrast to Hyena and StripedHyena. Architecture details of models trained for our scaling law analysis provided in Table S1.

#### Transformer++

We use a modern decoder-only Transformer architecture with rotary position embeddings (Su et al., 2024), pre-norm with root mean square layer normalization, and SwiGLU as channel mixer. The inner width of the SwiGLU is 4/3 the model width. We experimented with grouped-query attention (GQA) (Ainslie et al., 2023) and found minimal differences in final loss, suggesting the technique may be suited to DNA sequence modeling, in order to further reduce memory footprint during inference. All scaling results with Transformer++ do not use GQA.

#### Hyena

The Hyena baseline is designed with the same architecture improvements applied to the Transformer++ model. We replace all multi-headed self-attention layers with hyena layers, and use a modal canonical parametrization for the long convolution, with state dimension 8.

#### Mamba

We use the implementation of Mamba as provided by the authors in the public repository.

### B.5. Generating DNA sequences with Evo

We sample sequences from Evo using standard top-*k* and temperature-based methods for autoregressive models. Evo benefits from the fast recurrent mode of hyena layers, enabling lower latency and memory cost (Massaroli et al., 2024; Poli et al., 2023b). In particular, we use the recurrent form of the modal canonical form as shown in (Massaroli et al., 2024), first processing the prompt with a Fast Fourier Transform modified to return output and state. We use a cache for the states of short convolutions. Evo can generate sequences of up to 650k nucleotides on a single 80GB GPU, in contrast to other long context methods for dense Transformers requiring a larger number of nodes. We use standard kv-caching for rotary attention layers in StripedHyena.

#### Controllable generation

We follow standard language model prompting techniques that condition generation on a given prefix. For class-conditional generation we prompt with a single token, representing the desired class, or genomic sequence type (e.g. Cas system, IS200/605). The model can also be steered by prompting on desired DNA subsequences.

### B.6. Multimodal evaluations

#### B.6.1. Protein function prediction

We used DMS datasets to benchmark protein and nucleotide language models in their ability to predict mutational effects on protein function. In all cases, we used the nucleotide sequences reported by the original study authors. We limited our analysis to *E. coli* and human proteins, where *E. coli* protein information is contained in the Evo training dataset but where human proteins are not.

To compile the nucleotide information from *E. coli* DMS studies, we used all of the datasets listed in the ProteinGym benchmark for which we could also find nucleotide-level information reported by the original study authors. This resulted in six studies: a β-lactamase DMS by Firnberg et al. (2014), a β-lactamase DMS by Jacquier et al. (2013), a CcdB DMS by Adkar et al. (2012), a multi-protein thermostability dataset by Tsuboyama et al. (2023), an IF-1 DMS by Kelsic et al. (2016), and an Rnc DMS by Weeks and Ostermeier (2023).

To compile the nucleotide information from human DMS studies, we narrowed the scope of the set of datasets used in our human benchmark to the human datasets used by Livesey and Marsh (2023) to benchmark mutational effect predictors. We also limited our analysis to studies where we could also find nucleotide-level information reported by the original study authors. This resulted in six studies: a CBS DMS by Sun et al. (2020), a GDI1 DMS by Silverstein et al. (2022), a PDE3A DMS by Garvie et al. (2021), a P53 DMS by Kotler et al. (2018), a P53 DMS by Giacomelli et al. (2018), and a BRCA1 DMS by Findlay et al. (2018).

We compared Evo (pretrained with 8k context) to three nucleotide language models: GenSLM 2.5B, which was trained with a codon vocabulary on sets of genes from prokaryotic organisms (Zvyagin et al., 2023); Nucleotide Transformer 2B5_multi_species, which was trained with a 6-mer nucleotide vocabulary on genome sequences from prokaryotic and eukaryotic species (Dalla-Torre et al., 2023); and RNA-FM, which was trained on a single-nucleotide vocabulary on short ncRNA sequences (Chen et al., 2022). We also compared Evo to several protein language models trained on non-redundant, generic corpuses of protein sequences: CARP 640M (Yang et al., 2024), ESM-1v (Meier et al., 2021), ESM-2 650M, ESM-2 3B (Lin et al., 2023), ProGen2 large, and ProGen 2 xlarge (Madani et al., 2023). For studies that provide models with multiple parameter sizes, we selected the largest size on which we could perform inference with an 80 GB Nvidia H100 GPU on sequences from all of our benchmarked studies without exceeding GPU memory. We also included ESM-2 650M and ProGen2 large given that these models have sometimes shown better performance at function prediction than larger variants of these models (Notin et al., 2023).

To compare nucleotide and protein language models, we used all unique nucleotide sequences and their corresponding fitness values as reported by the original studies. Occasionally, we observed that the fitness values reported for nucleotide sequences differed from fitness values reported for protein sequences; in such cases, we used the fitness values reported for nucleotide sequences and evaluated the protein language models using the translated sequence. In cases where there are multiple nucleotide sequences for a single protein sequence due to different codon usage, the nucleotide language models were evaluated on each unique nucleotide sequence and the protein language models were evaluated on the coding sequence corresponding to each unique nucleotide sequence; this means that a protein language model could have been evaluated on the same protein sequence multiple times for a given study. Some studies report fitness values for mutations that involve stop codons; in such cases, we evaluated the nucleotide language model on the sequence containing the stop codon and excluded these examples from the protein language model benchmark.

We computed the Spearman correlation between the experimental fitness values and the sequence likelihood (for autoregressive language models) or the sequence pseudolikelihood (for masked language models). We assessed statistical significance of the Spearman correlation coefficient under a null hypothesis that the correlation coefficient is drawn from a *t*-distribution with *N* − 2 degrees of freedom, where *N* is the number of samples over which we compute the correlation. We used this null distribution to compute a *P* value based on the observed correlation. We used the scipy Python library (https://scipy.org/) to compute these values.

#### B.6.2. ncRNA function prediction

We used DMS datasets to benchmark protein and nucleotide language models based on their ability to predict mutational effects on ncRNA function. Given that no well established benchmark datasets exist for ncRNAs function prediction, we curated the literature for examples of ncRNA mutational scanning experiments. We obtained the following datasets: a ribozyme DMS by Kobori et al. (2015), a ribozyme DMS by Andreasson et al. (2020), a tRNA DMS by Domingo et al. (2018), a tRNA DMS by Guy et al. (2014), a ribozyme DMS by Hayden et al. (2011), a ribozyme DMS by Pitt and Ferré-D’Amaré (2010), and a rRNA mutagenesis study by Zhang et al. (2009).

We compared Evo (pretrained with 8k context) to the nucleotide language models described above. Similar to the methods applied to protein coding sequences above, we compiled experimental fitness values for each ncRNA variant. We computed the Spearman correlation between the experimental fitness values and the sequence likelihood (for autoregressive language models) or the sequence pseudolikelihood (for masked language models). Correlation coefficients and associate *P* values were computed as described above.

#### B.6.3. Gene expression prediction from regulatory DNA

To evaluate the model’s ability to learn properties of regulatory DNA, we used a dataset reported by Kosuri, et al. (2013) in which a set of *E. coli* promoters and a set of *E. coli* RBSs were combinatorially paired and the promoter-RBS pairs were experimentally tested for their effect on downstream mRNA and protein expression. We computed the sequence likelihood (for autoregressive language models) or the sequence pseudolikelihood (for masked language models) for each promoter-RBS pair, where we concatenated the sequence of the promoter directly with the sequence of the RBS.

We computed these likelihoods using Evo (pretrained with 8k context) and the three other nucleotide language models described above. We used these likelihoods to predict continuous mRNA expression values and binarized protein expression values as reported in the original study. Protein expression was binarized using a cutoff at which expression values above 100,000 were treated as positive and values below 100,000 were treated as negative, where this cutoff was based on the bimodal distribution of protein expression values reported in the original study. We used the Spearman correlation coefficient to quantify the predictive performance for mRNA expression and the AUROC to quantify predictive performance for protein expression. We assessed statistical significance of the Spearman correlation coefficient with a *t*-distributed *P*-value as described above. We assessed the statistical significance of the AUROC with a permutation-based method in which a null distribution is constructed by permuting the binary labels and recomputing the subsequence AUROC. We performed 100,000 permutations to construct this null distribution.

We also attempted to quantify how well a given promoter-RBS pair was represented in the Evo pretraining data, as we hypothesized that promoter-RBS pairs that are seen more often in nature, and would thereby have higher language model likelihood, are also pairs that are more likely to lead to higher gene expression. We attempted to align the promoter-RBS sequences to bacterial genomic sequences using three methods. First, we constructed a BLAST database over the full GTDB using the makeblastdb command with default parameters (Ye et al., 2006). We then used the blastn command with default parameters where for each promoter-RBS pair we queried the database for significant hits. We used the number of returned BLAST hits to score each promoter-RBS pair (having no hits was scored as zero). Second, we used mmseqs to construct databases over the set of promoter-RBS pairs and over the full GTDB using the mmseqs createdb command (––dbtype 2) to create nucleotide databases. We then used mmseqs createindex (––search-type 3) to create a nucleotide search index. We then conducted an all-by-all search (––cov-mode 2) to search for sequences in GTDB that aligned to the promoter-RBS queries. We used the number of significant alignments to score each promoter-RBS pair (having no alignments was scored as zero). Third, we attempted to align promoter-RBS sequences to the *E. coli* reference genome (RefSeq: GCF_000005845.2) using bowtie2. We used bowtie2-build with default parameters to build an index over the *E. coli* genome. We then treated promoter-RBS pairs as unpaired reads in a FASTA file and enabled multimapping with the -a flag to bowtie2. We used the number of alignments to score each promoter-RBS pair (having no alignments was scored as zero). We used the Spearman correlation coefficient to quantify predictive performance for mRNA expression and the AUROC to quantify predictive performance for protein expression and report the highest correlation values across the three attempted methods.

### B.7. CRISPR-Cas finetuning, generation, and downstream analysis

To generate CRISPR-Cas systems, we finetuned Evo by continuing to train the 8k-context pretrained model on a dataset of CRISPR-Cas sequences, which was curated as described above. We retained most of the hyperparameters used during pretraining but set the batch size to 524k tokens and an initial learning rate of 0.00009698, which was the learning rate at the final step of pretraining. During pretraining, we prepended a single class token corresponding to the type of Cas protein (Cas9, Cas12, or Cas13), which was identified as described above; this class token was then followed by the nucleotide sequence. We also modified the dataloader such that each sample provided to the model during training would begin with the first token of the CRISPR-Cas sequence and, if a sequence was shorter than the context length, we padded the sequence to the remaining context (where padding did not contribute to the loss computation). This ensured that each training sample would correspond to a single CRISPR-Cas sequence. We finetuned the model for approximately 10 epochs.

We prompted the model with a given class token for each sequence generation. We performed standard temperature-based and top-*k* autoregressive sampling (Chang and Bergen, 2023). In our generations, we performed an exhaustive sweep consisting of temperatures of 0.1, 0.3, 0.5, 0.7, 0.9, 1.0, and 1.3, and top-*k* values of 2 and 4. All sampled sequences were then combined into a single file and used for downstream extraction and analysis of candidate CRISPR systems.

The in silico Cas evaluation pipeline consisted of an initial open reading frame (ORF) search using Prodigal (Hyatt et al., 2010) and subsequent profiling of the extracted ORFs using hidden markov model (HMM) profiles for each Cas subtype. Sampled sequences with a positive pHMM hit with an E-value under 1 × 10^−3^ and a sequence length above a given threshold were further analyzed using the MinCED package to identify possible CRISPR arrays (Bland et al., 2007). Generations containing both a Cas ORF and a CRISPR array were then clustered using MMSeqs2 at a sequence identity of 90% and minimum coverage length of 75% (Steinegger and Söding, 2017). Finally, representative sequences from the clustering analysis were aligned against Cas ORF sequences in the training data with MMSeqs2 to quantify divergence from the training dataset.

Candidate sequences were selected from the cluster representatives within various sequence identity deciles and processed using AlphaFold2 to manually inspect structural similarities between generations and a crystal structure of wild-type SpCas9. Predicted Cas9 structures were aligned to SpCas9 (PDB: 4OO8) and its gRNA complex with ChimeraX and top candidates were chosen for further analysis. Possible recognition (REC) and nuclease (NUC) lobes of the sampled Cas9 structures were labeled by performing a multiple sequence alignment with the protein sequence of SpCas9. The MSA boundaries of the lobes in SpCas9 were used as boundaries for labeling the REC and NUC lobes in the sampled sequences. CRISPRtracrRNA was used to extract potential tracrRNA sequences from candidate generations and co-folded with the extracted crRNA sequence using RNAmultifold (Mitrofanov et al., 2022; Lorenz et al., 2011). Different combinations of tracrRNA and crRNA lengths were assessed as the resulting mature crRNA and tracrRNA sequences are not readily apparent from raw sequence data.

### B.8. IS200/IS605 finetuning, generation, and downstream analysis

To generate IS605 systems, we finetuned Evo by continuing to train the 8k-context pretrained model on a dataset of IS200/IS605 sequences, which was curated as described above. We retained most of the hyperparameters used during pretraining but set the batch size to 524k tokens and an initial learning rate of 0.00009698, which was the learning rate at the final step of pretraining. During pretraining, we prepended a generic start token to each sequence. We also modified the data loader such that each sample provided to the model during training would begin with the first token of the IS200/IS605 sequence and, if a sequence was shorter than the context length, we padded the sequence to the remaining context (where padding did not contribute to the loss computation), similar to the strategy described for CRISPR-Cas9 systems above. We finetuned the model for approximately 10 epochs.

We prompted the model with the start token for each sequence generation. We performed standard temperature-based and top-*k* autoregressive sampling (Chang and Bergen, 2023). In our generations, we performed an exhaustive sweep consisting of temperatures of 0.1, 0.3, 0.5, 0.7, 0.9, 1.0, and 1.3, and top-*k* values of 2 and 4. We sampled a total of 1,004,850 sequences.

We analyzed generated sequences using prodigal to identify coding sequences and proteins (Hyatt et al., 2010), followed by hmmsearch (-Z 1000000) using pHMMs to identify TnpA and TnpB sequences (Finn et al., 2011), and cmsearch (-Z 4) using covariance models developed in a previous publication (Meers et al., 2023) to identify candidate ωRNAs (Nawrocki and Eddy, 2013). Candidate TnpA sequences were kept if they had an E-value < 1 × 10−3 to the pHMM and if they covered at least 50% of the pHMM. Candidate TnpB sequences were kept if they had an E-value < 1 × 10−3 to at least one pHMM, if they covered at least 50% of the pHMM, and if they were between 300 and 600 amino acids in length.

Predicted TnpA and TnpB protein sequences were aligned back to proteins in the training set using MM-seqs2 (Steinegger and Söding, 2017). The top hit for each protein was extracted and separately aligned using the MAFFT-G-INS-I algorithm to estimate the amino acid identity across the full lengths of the two sequences (Katoh et al., 2002). To account for different start codons and to generate a more conservative percentage identity, these alignments were trimmed to the middle 80% of each sequence, end gaps were trimmed, and the amino acid identity was recalculated.

For loci that contained both a TnpA and a TnpB coding sequence, we used ESMFold (Lin et al., 2023) to predict atomic-level structures for each protein sequence. We reported the mean backbone atom pLDDT as a measurement of ESMFold prediction confidence. Example TnpA and TnpB proteins were aligned to the 2EC2 and 8BF8 PDB structures, respectively, using the cealign algorithm in PyMOL (Schrödinger, LLC, 2015). RNAfold from the ViennaRNA package was used to fold the predicted ωRNA with parameters “-d3-P rna_langdon2018.par” (Gruber et al., 2008; Langdon et al., 2018). We also used Evo to calculate the entropy of the conditional probabilities at each position in a given sequence. For example, the entropy at position *i* was calculated using the likelihoods *P*(*xi* |*x*1, …, *xi*−1) over the entire vocabulary. We then visualized these entropies alongside the annotated sequence positions for several canonical IS200/IS605 systems.

### B.9. Gene essentiality prediction

We obtained binary genome-wide essentiality results for 56 bacterial genomes from the DEG database (Zhang, 2004) in which coding genes are labeled with “essential” or “nonessential” binary labels. We also obtained genome-wide essentiality results for two phage genomes, lambda and P1, from Piya et al. (Piya et al., 2023) and used the binary labels assigned by the study authors based on the results of their CRISPRi screen.

To perform the in silico gene essentiality screen, we obtained the whole bacterial genome using the RefSeq IDs provided by DEG. We used RefSeq: NC_001416as the reference genome for lambda phage and RefSeq: NC_005856 as the reference genome for P1 phage. We iterated over all genes annotated as protein coding and computed a score with a nucleotide language model for each gene. To compute the score, we provided the language model with different levels of context: (1) the sequence of the gene only, (2) the sequence of the gene plus equally distributed context on both sides of the gene up to a total 8,192bp, or (3) the sequence of the gene plus equally distributed context on both sides of the gene up to a total 65,536 bp. If a gene extended beyond 8,192 bp, we used the first 8,192 bp of the gene sequences. We computed the score as the difference in log-likelihoods between a mutated sequence and the unmutated wildtype sequence. To mutate the sequence, we inserted multiple stop codons “TAATAATAATAGTGA” 12 nucleotides into the sequence; for the 8,192 and 65,536 bp context settings, we add context to both sides of the gene after the insertion. Additionally, for the 8,192 bp setting, we tested two other strategies: (1) inserting a single stop codon “TAA” 12 nucleotides into the sequence and (2) deleting the entire gene sequence (after which we provided 8,192 context centered on the deleted gene) (Figure S9). As an additional control, we also used the gene’s linear position in the reference genome as the value with which to predict essentiality. If a model were simply using positional information to make essentiality predictions, the performance would be similar to this control.

We used the change in log-likelihoods to predict the binary gene essentiality labels and compute the strength of the prediction with the AUROC score. We assessed the statistical significance of the AUROC with a permutation-based method in which a null distribution is constructed by permuting the binary labels and recomputing the subsequence AUROC. We performed 100,000 permutations to construct this null distribution.

### B.10. Genome-scale generation and evaluation

We used Evo pretrained at 131k context to sample twenty sequences up to lengths ∼650 kb. We sampled with a temperature of 0.7 and a top-*k* value of 4 following a standard autoregressive sampling procedure (Chang and Bergen, 2023). We prompted the model with four species-specific prompts, which were introduced during 131k pretraining, corresponding to the bacterial species *Mycoplasma genitalium*, *Staphylococcus aureus*, *Klebsiella pneumoniae*, and *Escherichia coli*. These prompts follow Greengenes-style lineage strings, which concatenate all taxa starting with the most ancestral and ending with the most current, separated by semicolons. A single character prefix is also added to each taxon indicating its rank. We sampled five sequences for each prompt, leading to a total of twenty sequences.

We evaluated these generations with CheckM (Parks et al., 2015), a tool that computes basic genome quality metrics based on whether a given long DNA sequence has similar properties as known bacterial genomes. CheckM uses Prodigal (Hyatt et al., 2010) to identify coding sequences and computes the coding density as one metric of genome quality. CheckM will also search for the presence of genes that are highly conserved across much of prokaryotic diversity. We divided all of our generations into five discrete segments of up to 131,072 bp (a total of 100 sequences) and computed the distribution of CheckM coding densities across these crops. As a positive control, we randomly selected 100 bacterial genomes from GTDB and used CheckM to compute the coding densities for 131,072 bp crops from these genomes. As a negative control, we generated 1,000 sequences of length 131,072 in which the four DNA base pairs were sampled uniformly at random. We then used CheckM to compute the coding densities on this random sequence. We also used tRNAscan-SE to search for tRNA sequences in our generated sequences and we used barrnap to search for rRNA sequences.

We used ESMFold to obtain atomic-level structure predictions for all of the Prodigal-defined coding sequences in each of our generations. We limited ESMFold structure predictions to coding sequences between 100 and 1024 amino acids, inclusive. We computed the mean backbone pLDDT for all predicted structures. We used the biotite Python package to compute the percentages of secondary structure elements for all predicted structures. We used FoldSeek easy-search to perform efficient TM-based alignment (––alignment-type 1), and all other parameters set to their default values, to perform an all-by-all structural search between ESMFold structures corresponding to Evo-generated sequences and the structure predictions for UniRef50 provided in the AlphaFold Protein Structure Database (https://alphafold.ebi.ac.uk/). Structure alignments were scored as the average of the query TMscore and the target TMscore, where a score greater than 0.4 was considered a structural match. We used these structural matches, along with GO terms assigned to UniRef50 clusters, to infer GO terms for the Evo-generated proteins as well. We used PyMOL to visualize protein structures corresponding to the five GO “molecular function” terms with the most representation among the Evo generated proteins.

## C. Supplementary tables and figures

**Figure S1.**
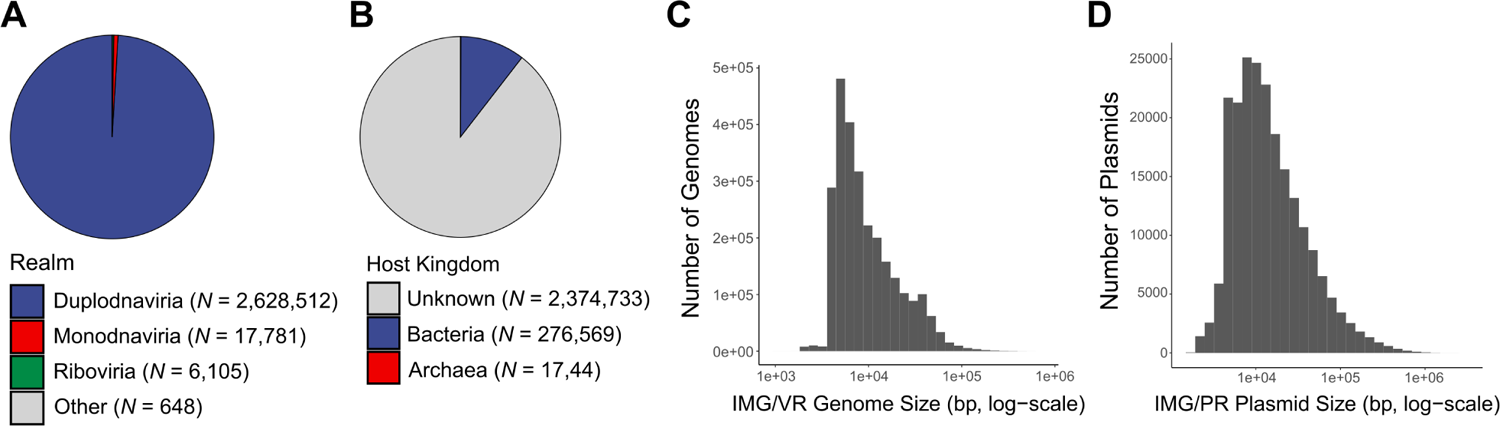
Pretraining data. Statistics of IMG/VR and IMG/PR. (**A**) A pie chart depicting the composition of viral realms in the IMG/VR subset of the pretraining dataset. (**B**) A pie chart depicting the composition of host kingdoms in the IMG/VR subset of the pretraining dataset. We excluded viruses that are likely to infect eukaryotic hosts (**Methods**). (**C**) The distribution of sequence lengths in the IMG/VR subset of the pretraining dataset. (**D**) The distribution of sequence lengths in the IMG/PR subset of the pretraining dataset.

**Figure S2.**
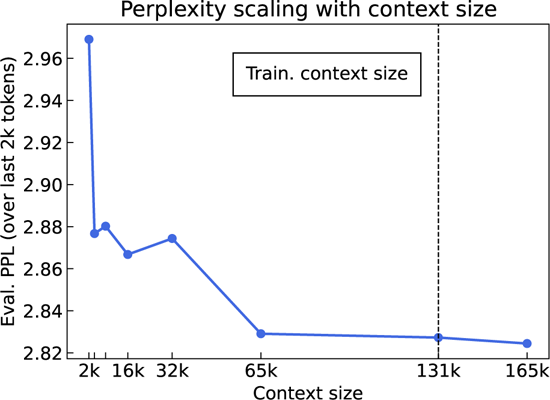
Perplexity scaling in context length. Perplexity on a subset of the OpenGenome validation set with Evo 131k as a function of sequence length, or context length. The perplexity is computed over the last 2048 nucleotides of each sequence, with increasing lengths of the prefix and thus of the context available to the model. We observe perplexity to continually decrease beyond the training context length at 131k, indicated by the vertical dashed line.

**Figure S3.**
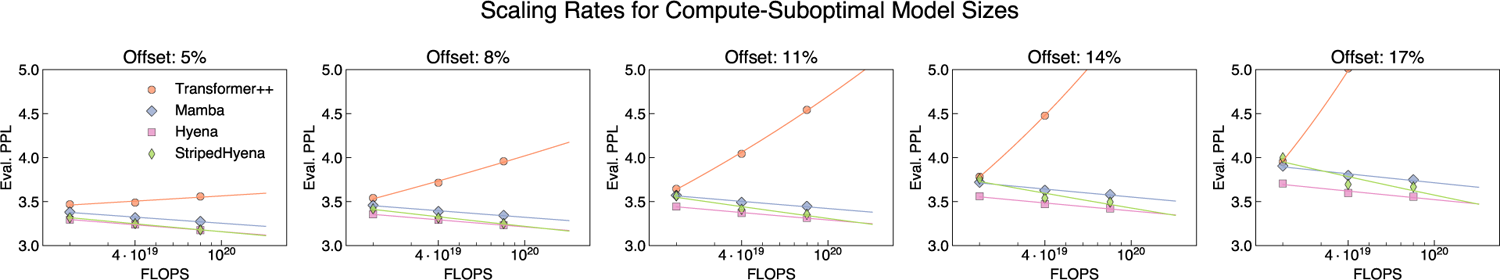
Scaling rates for compute-suboptimal model sizes by architecture. We quantify the effect on perplexity scaling caused by a suboptimal allocation of compute budget to model or dataset size (e.g., training a smaller model for more tokens). We estimate the compute-optimal model size (Figure S5) for each compute budget, then reduce it by a percentage (the *offset*). The corresponding perplexity is obtained via the IsoFLOP curves (Figure 1F). Transformer++ perplexity scaling rapidly degrades outside the compute-optimal frontier, in contrast to Hyena and StripedHyena.

**Figure S4.**
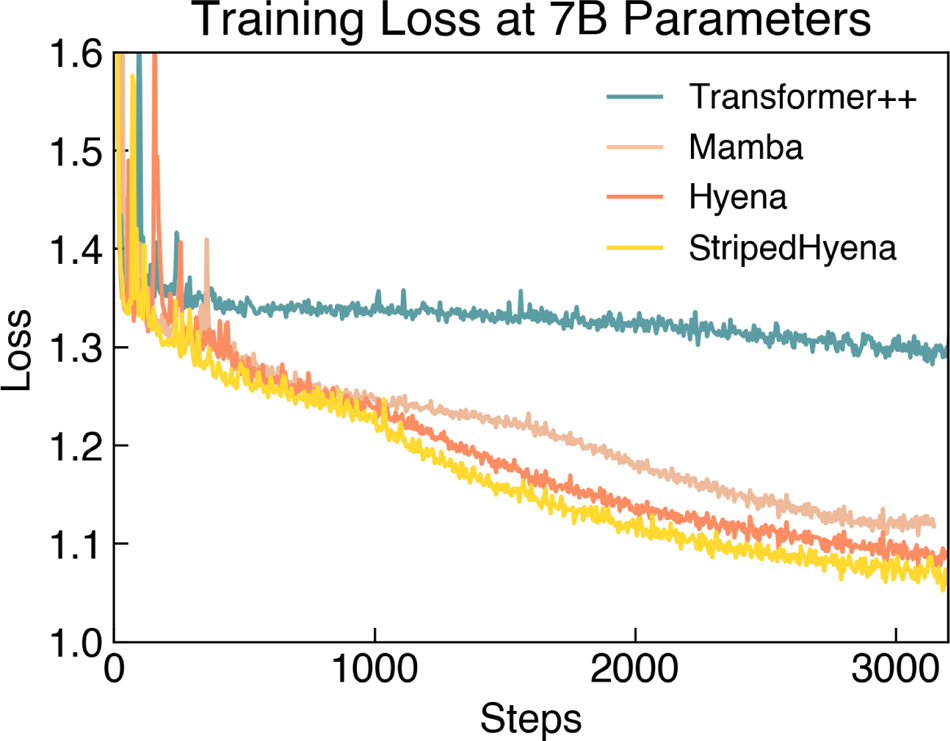
Direct comparison of training loss curves during hyperparameter tuning for 7B models. We tune several Transformer++ and Mamba models as baselines, sweeping learning rates, batch size, sequence lengths, and model depth vs. width ratio. In all settings, Hyena and StripedHyena outperformed Transformer++ and Mamba, where both baselines experienced instability during training.

**Figure S5.**
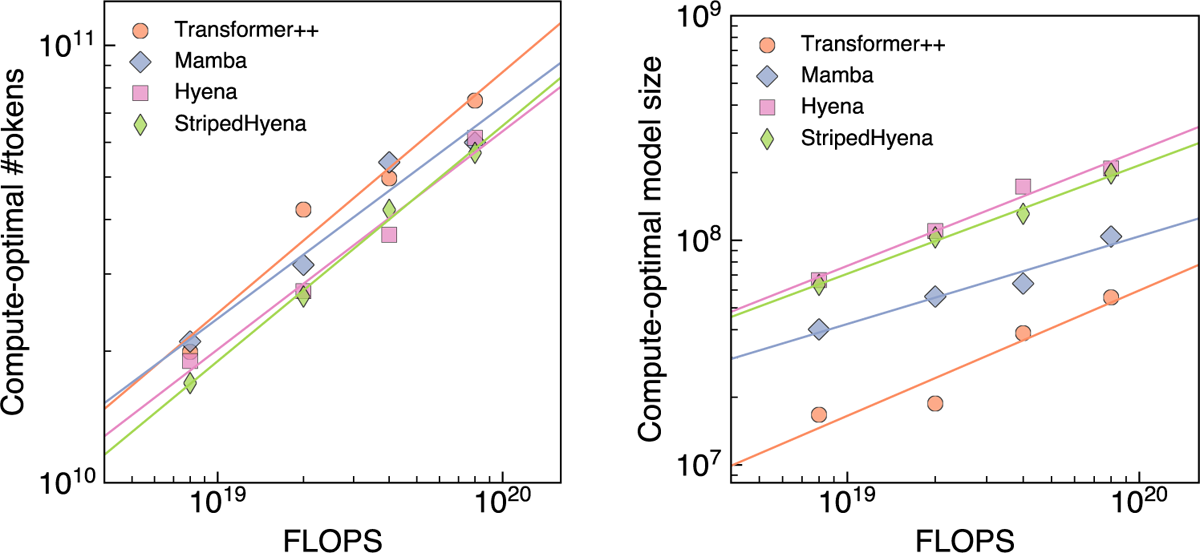
Compute-optimal tokens and model size by model. Compute-optimal allocation to dataset size (number of tokens) and model size (number of parameters) for each compute budget, measured in FLOPS.

**Figure S6.**
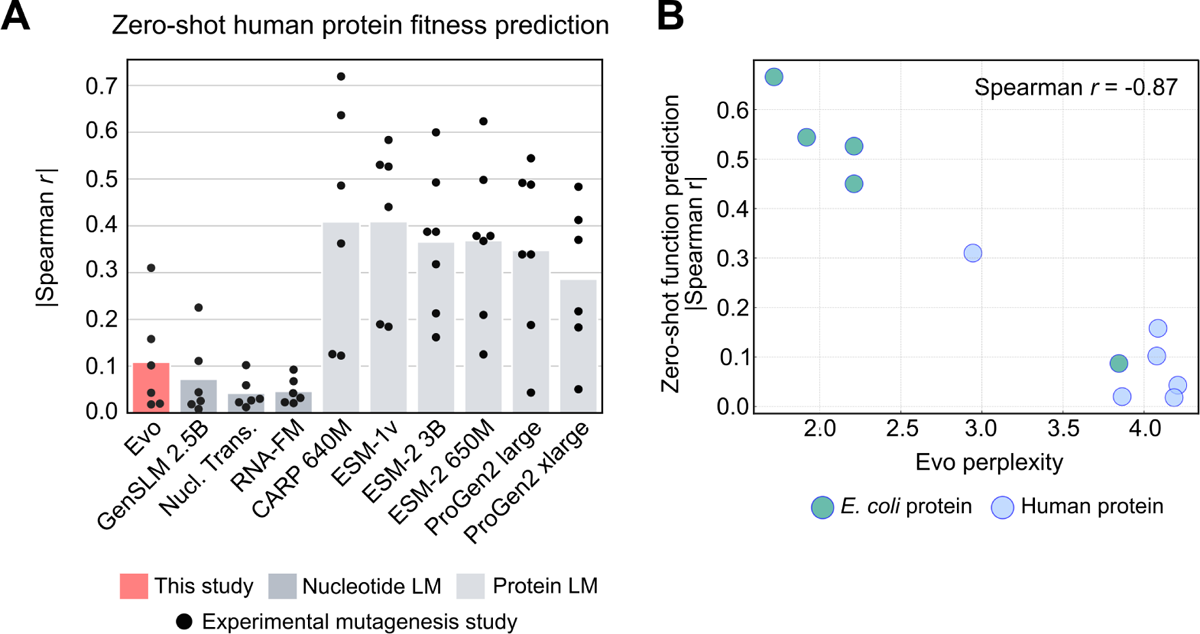
Performance of Evo on mutational effect prediction for human proteins. (**A**) Predictive performance of nucleotide and protein language models on mutational effect prediction for human proteins, measured via Spearman correlation. Bar height indicates the mean; each dot indicates a different DMS study. LM: language model; Nucl. Trans.: Nucleotide Transformer. Related to **Figure 2B**. (**B**) Relationship between the Evo perplexity of the wildtype nucleotide sequence (horizontal axis) and the ability for Evo to perform zero-shot mutational effect prediction for that protein as measured via Spearman correlation (vertical axis). Each dot corresponds to a different protein; dots are colored as to whether they are *E. coli* (teal) or human (blue) proteins. We observed a strong negative correlation (Spearman *r* = −0.87) between perplexity and zero-shot function prediction performance.

**Figure S7.**
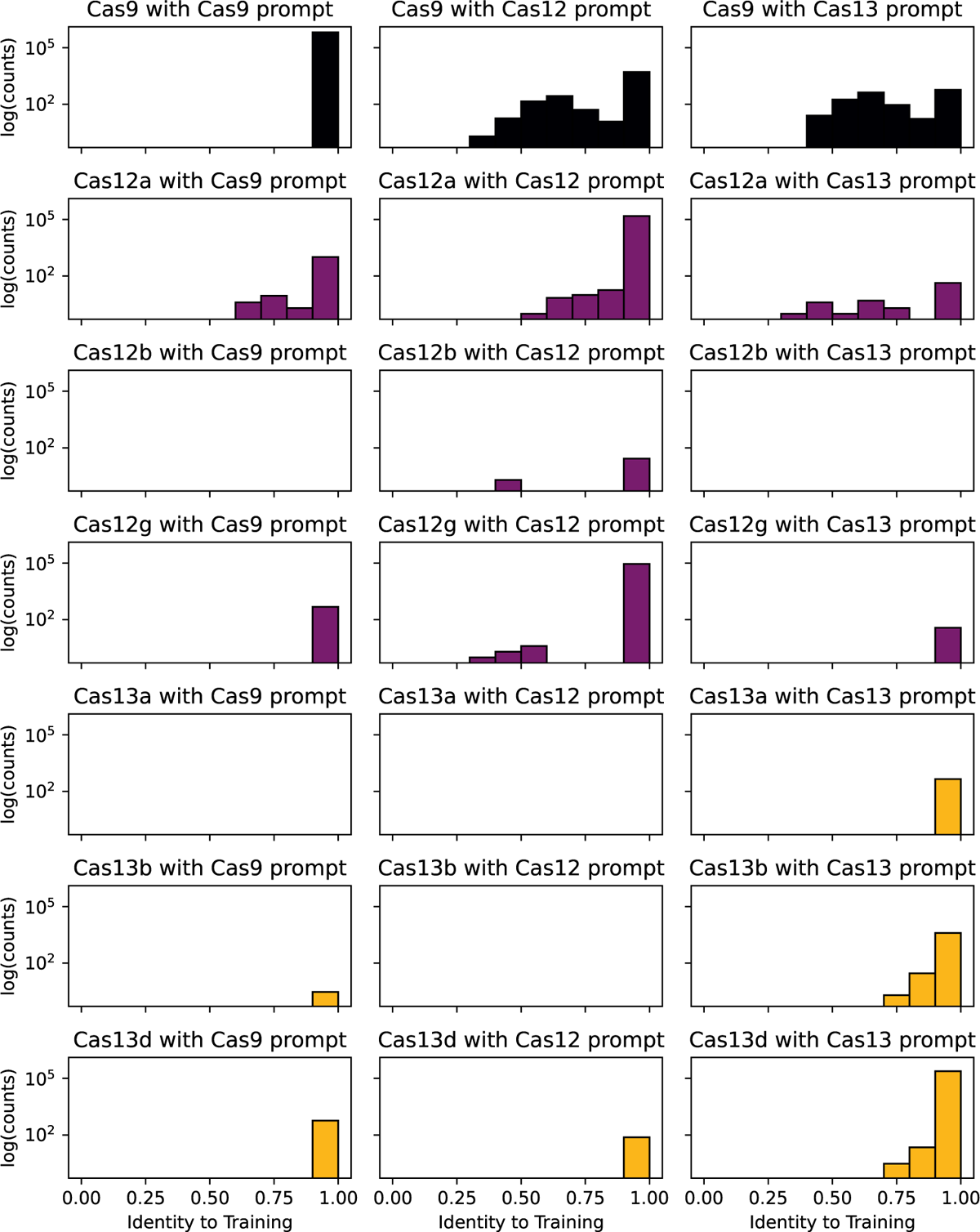
Breakdown of Cas generation diversity by prompt token. Evo can generate diverse Cas loci across subtypes through conventional prompting as well as through cross-type prompting. Evo was prompted with “cas9”, “cas12”, or “cas13” special tokens and each resulting set of generations was analyzed using pHMMs for each Class 2 Cas subtype and compared against training data to observe divergence from the training dataset.

**Figure S8.**
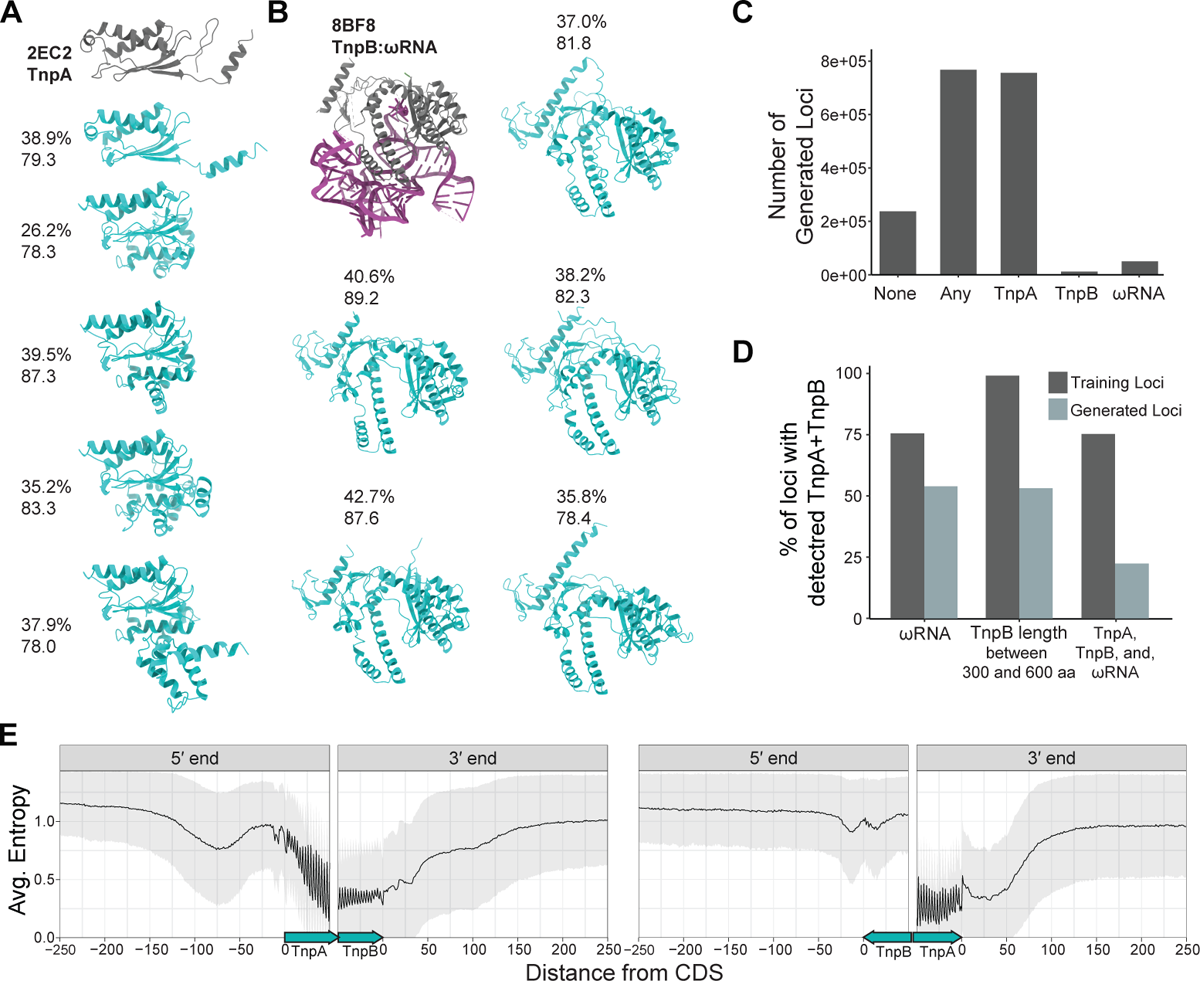
Additional analysis of generated IS200/IS605 sequences and the finetuned model. (**A**) Additional examples of diverse TnpA-like proteins with detected Y1 HuH domains and high mean backbone atom pLDDT. Labels indicate the percent amino acid identity to closest protein hit found in training set or NCBI nr and pLDDT. PDB structure 2EC2 shown for reference. (**B**) Examples of diverse generated TnpB proteins. PDB structure 8BF8 shown for reference. (**C**) A summary of the 1,004,850 sequences generated using the model finetuned on IS200/IS605 loci. Sequences have a detected TnpA, TnpB, or ωRNA (“Any”), a TnpA coding sequence (“TnpA”), a TnpB coding sequence (“TnpB”), or a ωRNA (“ωRNA”). (**D**) A comparison of the percentage in each category across the training set and generated sequences for sequences with a detected TnpA and TnpB coding sequence. Categories include sequences with a detected ωRNA (“ωRNA”); sequences encoding a TnpB protein between 300 and 600 aa in length; and sequences with a TnpA, a TnpB protein between 300 and 600 aa in length, and an ωRNA. (**E**) The average entropy within 250 nt of the 5^′^ and 3^′^ends of IS605 coding sequences, including 50 nt of the CDS itself. The entropy was calculated at each position across IS605 sequences in the training set (*N* = 10,419). Sequence positions were aligned with respect to the beginning and end of each respective CDS. (Left) All sequences with a TnpA followed by a TnpB. (Right) Sequences where the TnpB precedes the TnpA on the forward strand. Gray ribbon indicates the standard deviation of the entropy values.

**Figure S9.**
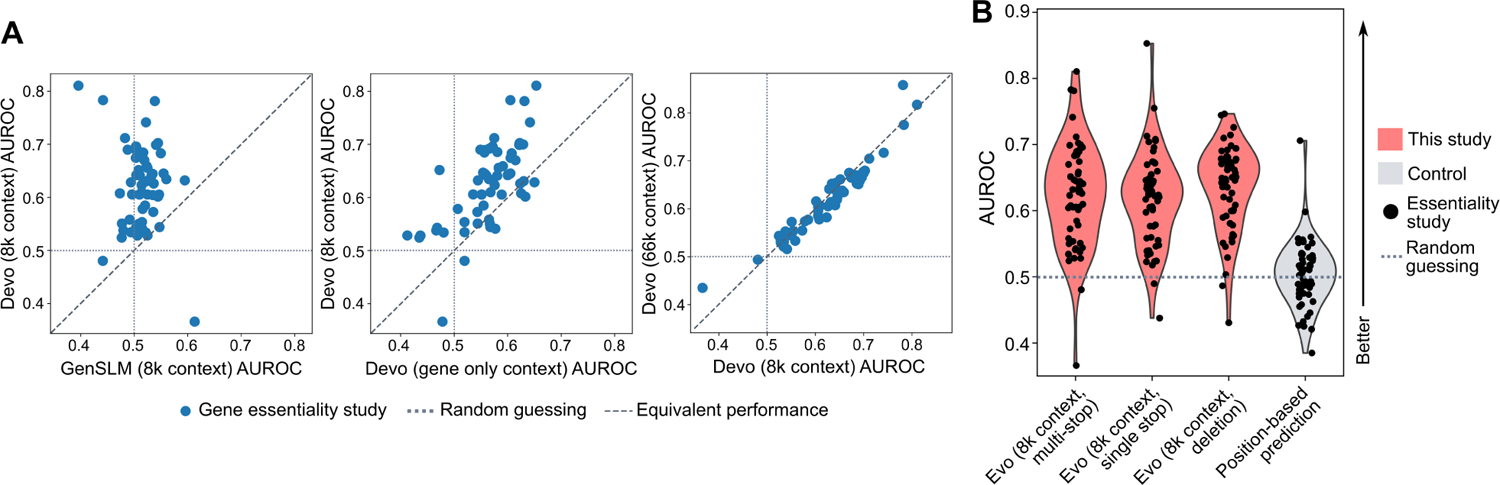
Gene essentiality prediction under different settings. (**A**) Scatterplots that compare the AUROC values between different models or different context windows. Axis labels are the same as the horizontal labels in **Figure 5C**. Each dot corresponds to a different whole-genome essentiality study. Related to **Figure 5C**. (**B**) Gene essentiality prediction performance for across 58 studies (each dot corresponds to a different study). We performed in silico mutagenesis of each coding sequence in a genome and commuted the change in Evo likelihood, which we used to predict gene essentiality. “Evo (8k context, multi-stop)” indicates a mutagenesis strategy that inserts multiple stop codons at the beginning of each coding sequence. “Evo (8k context, single stop)” indicates a mutagenesis that inserts a single stop codon at the beginning of each coding sequence. “Evo (8k context, deletion)” indicates a mutagenesis strategy that deletes the entire sequence of the gene. “Position-based prediction” indicates a prediction strategy (not using Evo) in which we use the position of a gene in the reference genome annotation as the predictor variable for gene essentiality. See **Methods** for more details. Related to **Figure 5C**.

**Figure S10.**
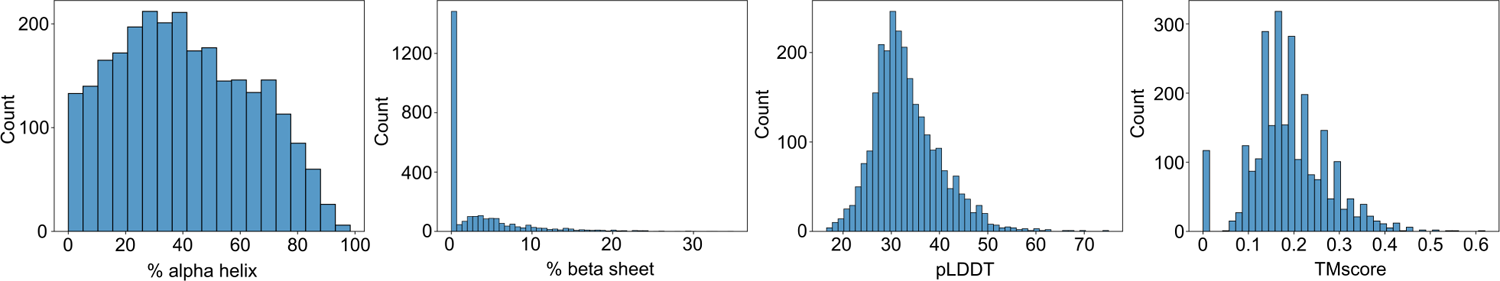
Statistics for ESMFold structure predictions of Evo-generated protein coding sequences. Histograms representing the distribution of statistics computed on ESMFold-predicted structures. These structures correspond to coding sequences found on five Evo-generated sequences, each of length ∼650 kb. These statistics are, from left to right: the percentage of residues in alpha helices, the percentage of residues in beta sheets, the mean backbone pLDDT, and the TMscore to the closest UniRef50 structure in the AlphaFold Protein Structure Database as determined by FoldSeek easy-search.

---

**Table S1.** Scaling laws model settings for Transformer++, Hyena, StripedHyena and Mamba. Layer number for Mamba is doubled (a single block corresponds to two Mamba layers and the dedicated channel mixer layer is removed, as described in (Gu and Dao, 2023)). Parameter counts vary slightly for each architecture.

| PARAMS (M) | D_MODEL | GLU_SIZE | KV_SIZE | N_HEADS | N_LAYERS | LEARNING RATE |
| --- | --- | --- | --- | --- | --- | --- |
| 1 | 128 | 336 | 64 | 2 | 4 | 9.77E-04 |
| 6 | 320 | 848 | 64 | 5 | 5 | 9.57E-04 |
| 17 | 448 | 1200 | 64 | 7 | 7 | 9.36E-04 |
| 29 | 512 | 1360 | 64 | 8 | 9 | 9.15E-04 |
| 40 | 576 | 1536 | 64 | 8 | 10 | 8.95E-04 |
| 59 | 640 | 1696 | 64 | 10 | 12 | 8.70E-04 |
| 69 | 640 | 1712 | 64 | 10 | 14 | 8.56E-04 |
| 84 | 704 | 1872 | 64 | 11 | 14 | 8.37E-04 |
| 99 | 768 | 2048 | 64 | 12 | 14 | 8.18E-04 |
| 114 | 768 | 2048 | 64 | 12 | 16 | 8.00E-04 |
| 121 | 768 | 2048 | 64 | 12 | 17 | 7.75E-04 |
| 135 | 768 | 2048 | 64 | 12 | 19 | 7.50E-04 |
| 158 | 832 | 2224 | 64 | 13 | 19 | 7.25E-04 |
| 175 | 832 | 2224 | 64 | 13 | 21 | 7.00E-04 |
| 203 | 896 | 2384 | 64 | 14 | 21 | 6.75E-04 |
| 232 | 896 | 2384 | 64 | 14 | 24 | 6.50E-04 |
| 266 | 960 | 2560 | 64 | 15 | 24 | 6.25E-04 |
| 303 | 1024 | 2736 | 64 | 16 | 24 | 6.00E-04 |
| 383 | 1152 | 3072 | 64 | 18 | 24 | 5.66E-04 |
| 473 | 1280 | 3408 | 64 | 20 | 24 | 5.33E-04 |
| 572 | 1408 | 3760 | 128 | 11 | 24 | 5.00E-04 |
| 680 | 1536 | 4096 | 128 | 12 | 24 | 4.75E-04 |
| 798 | 1664 | 4432 | 128 | 13 | 24 | 4.55E-04 |
| 926 | 1792 | 4784 | 128 | 14 | 24 | 4.33E-04 |
| 1063 | 1920 | 5120 | 128 | 15 | 24 | 4.15E-04 |
| 1209 | 1920 | 5120 | 128 | 15 | 25 | 4.11E-04 |

**Table S2.**
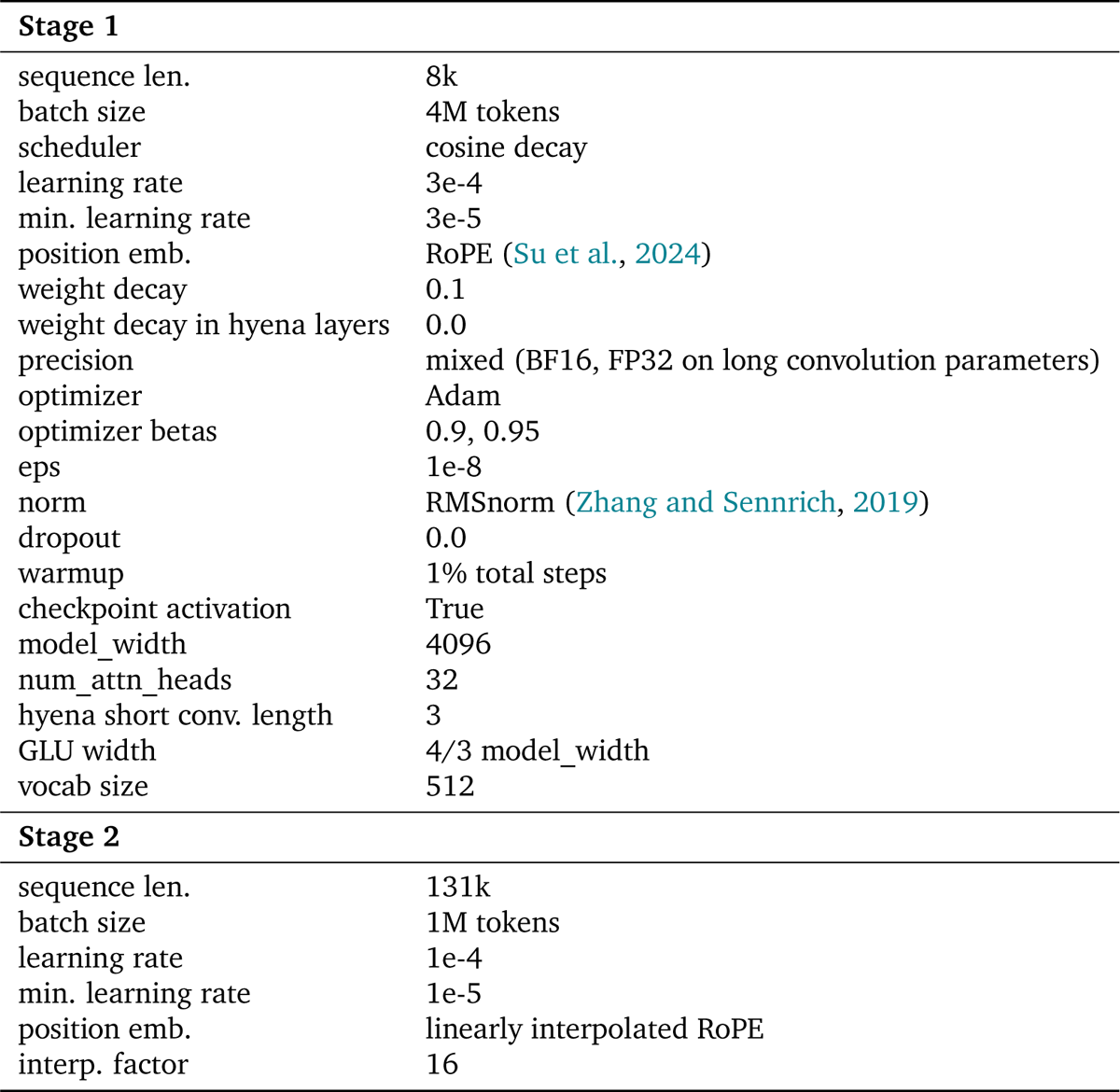
StripedHyena 7B hyperparameters settings at pretraining. Hyperparameters used for pretraining at 7B are shown by stage, where stage 2 highlights differences from stage 1.

| Stage 1 |  |
| --- | --- |
| sequence len. | 8k |
| batch size | 4M tokens |
| scheduler | cosine decay |
| learning rate | 3e-4 |
| min. learning rate | 3e-5 |
| position emb. | RoPE (Su et al., 2024) |
| weight decay | 0.1 |
| weight decay in hyena layers | 0.0 |
| precision | mixed (BF16, FP32 on long convolution parameters) |
| optimizer | Adam |
| optimizer betas | 0.9, 0.95 |
| eps | 1e-8 |
| norm | RMSnorm (Zhang and Sennrich, 2019) |
| dropout | 0.0 |
| warmup | 1% total steps |
| checkpoint activation | True |
| model_width | 4096 |
| num_attn_heads | 32 |
| hyena short conv. length | 3 |
| GLU width | 4/3 model_width |
| vocab size | 512 |
| Stage 2 |  |
| sequence len. | 131k |
| batch size | 1M tokens |
| learning rate | 1e-4 |
| min. learning rate | 1e-5 |
| position emb. | linearly interpolated RoPE |
| interp. factor | 16 |

**Table S3.**
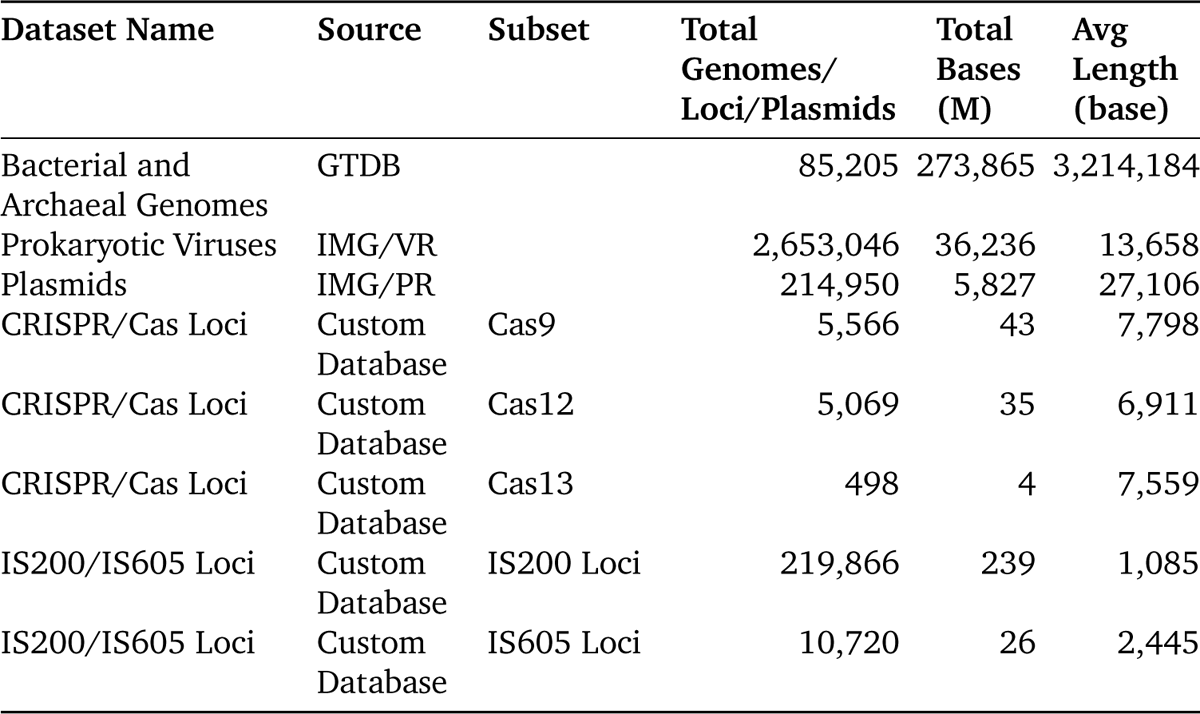
Summary statistics for the OpenGenome datasets. See B.2 for further details on the dataset sources and curating process.

| Dataset Name | Source | Subset | Total Genomes/<br>Loci/Plasmids | Total Bases<br>(M) | Avg Length<br>(base) |
| --- | --- | --- | --- | --- | --- |
| Bacterial and Archaeal Genomes | GTDB |  | 85,205 | 273,865 | 3,214,184 |
| Prokaryotic Viruses | IMG/VR |  | 2,653,046 | 36,236 | 13,658 |
| Plasmids | IMG/PR |  | 214,950 | 5,827 | 27,106 |
| CRISPR/Cas Loci | Custom Database | Cas9 | 5,566 | 43 | 7,798 |
| CRISPR/Cas Loci | Custom Database | Cas12 | 5,069 | 35 | 6,911 |
| CRISPR/Cas Loci | Custom Database | Cas13 | 498 | 4 | 7,559 |
| IS200/IS605 Loci | Custom Database | IS200 Loci | 219,866 | 239 | 1,085 |
| IS200/IS605 Loci | Custom Database | IS605 Loci | 10,720 | 26 | 2,445 |

